# Hyperactive GluN2B impairs neuroplasticity and cognition in phenylketonuria

**DOI:** 10.1101/2024.06.04.597492

**Authors:** Woo Seok Song, Young-Soo Bae, Sang Ho Yoon, Myoung-Hwan Kim

## Abstract

Phenylketonuria (PKU), an inborn error of phenylalanine (Phe) metabolism, is a common cause of intellectual disability. However, the mechanism by which elevated Phe levels causes cognitive impairment remains unclear. Here, we show that submillimolar Phe perturbs synaptic plasticity through the hyperactivation of GluN2B-containing NMDARs. L-Phe exhibited dose-dependent bidirectional effects on NMDA-induced currents, without affecting synaptic NMDARs in hippocampal CA1 neurons. L-Phe-induced hyperactivation of extrasynaptic GluN2B resulted in an activity-dependent downregulation of AMPARs during burst or sustained synaptic activity. Administration of L-Phe in mice decreased neural activity and impaired memory, which were blocked by pretreatment of GluN2B inhibitors. Furthermore, pharmacological and virus-mediated suppression of GluN2B reversed impaired learning in the Pah^Enu2^ PKU model. Collectively, these results suggest that the concentration of Phe in the cerebrospinal fluid of patients with PKU perturbs extrasynaptic NMDAR and synaptic plasticity, and that suppression of GluN2B may be a therapeutic strategy for improving cognitive function in patients with PKU.

## Introduction

Phenylketonuria (PKU) is caused by a deficiency in the activity of phenylalanine hydroxylase (Pah), which catalyzes the conversion of phenylalanine (Phe) to tyrosine (Tyr) in the liver^1,2^. Elevated Phe in the blood is transported across the blood-brain barrier (BBB) and causes severe intellectual disability [intelligence quotient (IQ) <30]. Treatment with restricted Phe intake (low-Phe diet) starting at an early age prevents severe intellectual disability. However, abnormal cognitive outcomes have been consistently observed in both continuously and early treated patients with PKU (_ETP_PKU)^3–7^.

Multiple studies have reported a high rate of white matter abnormalities (WMA), probably due to reduced cerebral protein synthesis^8^, in both untreated and _ETP_PKU ^9–12^. However, clinical studies have found no significant association between the extent of WMA and cognitive outcomes^10–14^. Instead, a clear correlation was observed between Phe levels and cognitive impairments in _ETP_PKU^3,4,13,15,16^.

Despite several decades of research that has enabled a biochemical understanding of the pathogenic mechanisms of PKU, one central unresolved question remains: How do elevated Phe levels cause cognitive impairments in patients with PKU? Previous studies have demonstrated that L-Phe at concentrations observed in or higher than the PKU serum reduces NMDAR- and AMPAR-mediated currents in cultured hippocampal and cortical neurons^17–19^. However, accumulating evidence from clinical and preclinical studies has revealed that the Phe concentration in the brain (Phe_brain_) or cerebrospinal fluid (Phe_CSF_) is substantially lower than that in the blood (Phe_blood_) in both patients with PKU and Pah^Enu^^2^ PKU mouse models^20–25^. Mean blood-brain ratios of Phe in _ETP_PKU were 4.0-4.12^21,24^, and that mean Phe_brain_ was ∼250 μmol/L or 270 μmol/kg^21,22,24^. Even in untreated children with classical PKU, the mean Phe_CSF_ was 399 μmol/L^20^. Notably, the oral Phe loading in _ETP_PKU increased the Phe_brain_ from 250 to 400 μmol/L and concomitantly shifted the dominant peak of the electroencephalogram (EEG) background activity to the lower-frequency spectrum^22^. These observations led us to hypothesize that submillimolar levels of Phe would affect neural circuits and brain activity in PKU.

To understand the neurophysiological mechanisms underlying cognitive impairments in PKU, we investigated the effects of submillimolar Phe on synaptic transmission and plasticity. Overall, we show that Phe in the PKU CSF hyperactivates GluN2B and perturbs synaptic plasticity via activity-dependent downregulation of AMPARs.

## Results

### Effects of L-Phe on NMDAR currents

The influence of Phe levels in PKU cerebrospinal fluid (CSF) on NMDAR-mediated synaptic transmission is unknown. Herein, we examined the effect of various concentrations of L-Phe on NMDAR-mediated excitatory postsynaptic currents (NMDAR-EPSCs) at the Schaffer collateral (SC)-CA1 synapse using hippocampal slices from adult mice. Unexpectedly, the peak amplitudes of NMDAR-EPSCs at this synapse were not affected by any of the tested doses (0.1 – 5 mM) of L-Phe (Fig. 1a and Supplementary Fig. 1). In contrast to synaptic transmission, however, L-Phe had a dose-dependent bidirectional effect on the current induced by NMDA perfusion (I_NMDA_) (Fig. 1b,c); L-Phe concentrations lower than 1 mM significantly enhanced the I_NMDA_ with the maximum effect at 250 μM, which is fairly close to the mean CSF concentration of L-Phe observed in patients with PKU^20–24^. In contrast, higher concentrations (5 and 10 mM) of L-Phe, similar to previous studies^17,18^, reduced I_NMDA_ significantly. Because GluN2A and GluN2B are the main types of NMDARs in the forebrain^26^, we examined the action of 250 μM L-Phe on each type of NMDAR. The enhancement in I_NMDA_ caused by L-Phe was sensitive to antagonists selective for GluN2B, but not for GluN2A-containing NMDARs (Fig. 1d,e).

**Fig. 1:**
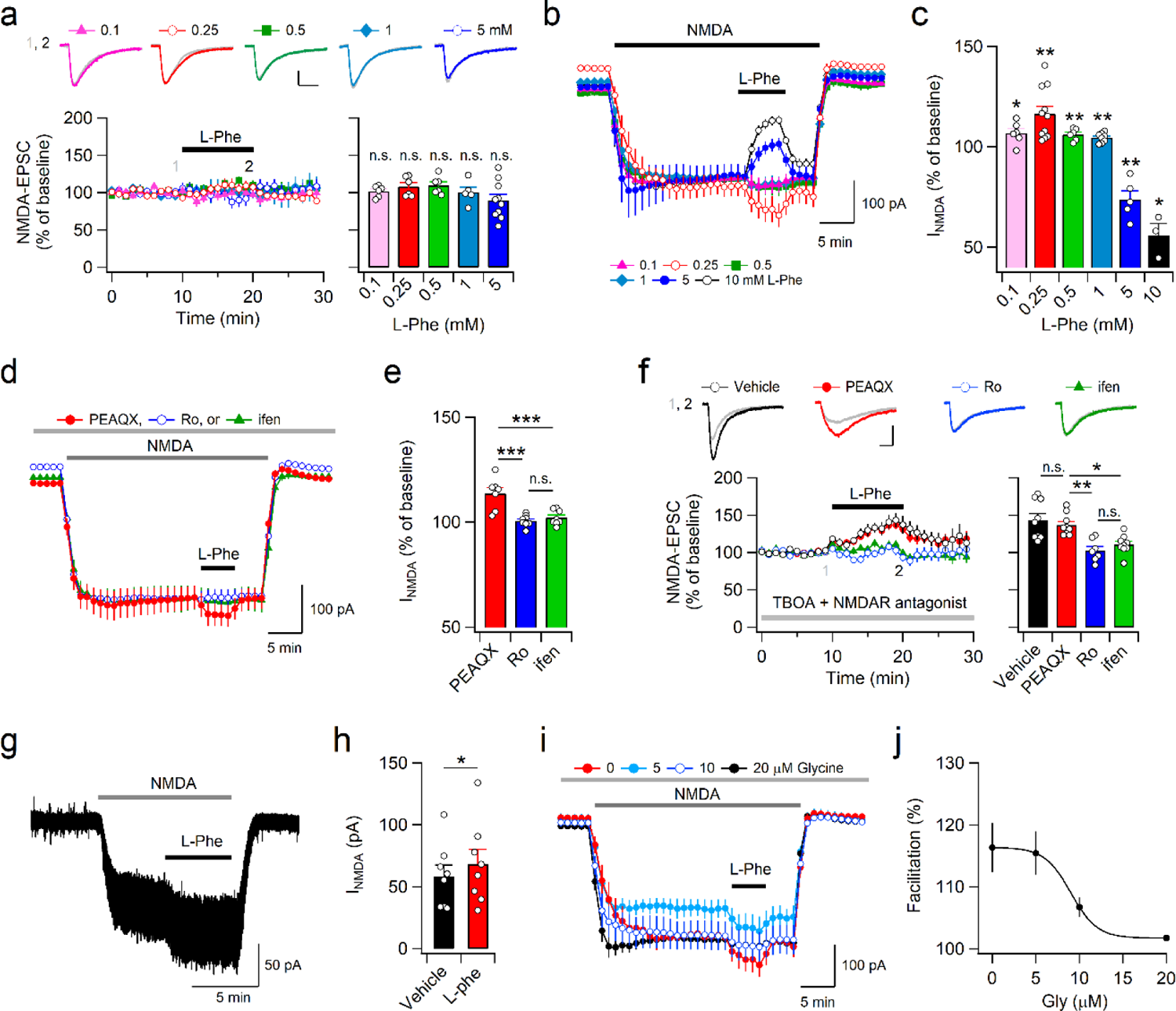
Submillimolar L-Phe increases the activity of GluN2B-NMDARs. **a**, Representative traces of NMDA-EPSCs (top) obtained at the indicated time points (1, 2), and the time course of the peak amplitudes (bottom, left) of NMDA-EPSCs measured with a holding potential of −40 mV in CA1 neurons. Scale bars, 50 ms and 50 pA. L-Phe had no effect on the peak amplitudes of NMDA-EPSCs at the SC-CA1 synapse (bottom, right). **b**, In the presence of NBQX and picrotoxin, bath application of NMDA (5−10 μM) induced an inward current in CA1 pyramidal neurons. Following a stable baseline of I_NMDA_ recording, different doses of L-Phe were perfused with NMDA for 5 min. **c**, L-Phe exhibits dose-dependent bidirectional effects on I_NMDA_ in CA1 pyramidal neurons. **d**, I_NMDA_ was induced by 3−12 μM NMDA in the presence of GluN2A or GluN2B blockers. **e**, L-Phe-induced facilitation of I_NMDA_ was blocked by Ro (2 μM) or ifen (6 μM) but not by PEAQX (0.5 μM). **f**, L-Phe (0.25 mM) increased NMDAR-EPSCs in the presence of TBOA (10 μM). Representative traces (top) and the time course of the peak amplitudes (bottom, left) of NMDAR-EPSCs. Scale bars, 50 ms and 50 pA. L-Phe induced facilitation of NMDAR-EPSCs in each condition (bottom, right). **g**, **h**, A sample trace (**g**) and summary (**h**) of I_NMDA_ measured before and during L-Phe perfusion in HEK293 cells expressing hGluN1.1a and hGluN2b. **i**, Addition of 5, 10, and 20 μM glycine attenuated L-Phe-induced I_NMDA_ facilitation. **g**, **i**, I_NMDA_ was induced by 30 (**g**) and 5 (**i**) μM NMDA. **j**, Concentration relationship between L-Phe-induced facilitation of I_NMDA_ and added glycine concentration in CA1 pyramidal cells.

We therefore wondered whether the absence of an effect of L-Phe on NMDAR-EPSCs stemmed from the subcellular location of NMDARs^27,28^. In the presence of the glutamate reuptake inhibitor DL-threo-β-Benzyloxyaspartic acid (TBOA), which promotes glutamate spillover to extrasynaptic areas^29^, L-Phe significantly enhanced NMDAR-EPSCs, indicating that L-Phe mainly influences the activity of extrasynaptic NMDARs at SC-CA1 synapses (Fig. 1f). Similar to I_NMDA_, the effect of L-Phe on NMDAR-EPSCs was still observed under GluN2A inhibition, but was blocked by the GluN2B inhibitors Ro25-6891 (Ro) and ifenprodil (ifen). The absence of an effect of L-Phe on the NMDAR-EPSCs is unlikely to stem from saturation of synaptic NMDARs^30^. In the absence of TBOA, the addition of D-serine, but not glycine^28^, to artificial cerebrospinal fluid (ACSF) significantly increased the amplitude of NMDAR-EPSCs (Supplementary Fig. 2). Moreover, L-Phe increased I_NMDA_ in HEK293 cells expressing the human GluN1.1a and GluN2b receptors (Fig. 1g,h), indicating a direct action of L-Phe on GluN2B-containing NMDARs. Previous studies have reported that L-Phe competes for the glycine-binding site in NMDARs and that attenuation of I_NMDA_ at high concentrations of L-Phe is dependent on the concentration of glycine^17,18^. Glycine exhibits an approximately 10-fold higher affinity for GluN2B-than for GluN2A-containing NMDARs^31^. As L-Phe induced facilitation in I_NMDA_ and NMDAR-EPSCs under TBOA treatment was observed at ambient glycine levels in the hippocampal slices, we tested whether the levels of glycine would affect the L-Phe-induced facilitation of I_NMDA_ in mouse hippocampal slices. Addition of glycine to ACSF decreased the magnitude of L-phe-induced I_NMDA_ facilitation in a dose-dependent manner (Fig. 1i,j). However, facilitation of I_NMDA_ was still observed up to 10 μM glycine, which is the normal concentration in the human CSF^32^. Considering that CSF glycine concentration was markedly reduced in untreated PKU infants, but was within the normal range in children and adults with PKU^33,34^, these results suggest that Phe levels in the PKU CSF likely facilitate the neurotransmission and signaling of GluN2B-containg NMDARs.

### L-Phe impairs synaptic plasticity

The neurological complications of PKU are attributable to altered synaptic function caused by long-term exposure to Phe and/or Phe in the CSF. We therefore examined whether the expression levels of the NMDAR subunits were altered in the brains of PKU mouse (Pah^Enu2^). In contrast to the changes in glutamate receptor expression observed in the BTBR strain^18,35^, adult PKU mouse with a C57BL6N background exhibited reduced GluN1 expression in the hippocampal homogenates but not in the P2 (synaptosome) fractions (Fig. 2a,b). GluN2A and GluN2B expression levels in both the total and P2 fractions of hippocampal homogenates did not differ between WT and PKU mice. Consistent with the normal expression levels of NMDAR subunits in the P2 fraction, the NMDA-AMPA ratios, including AMPAR-mediated synaptic transmission, at the SC-CA1 synapse in Pah^Enu2^ mice did not change (Fig. 2c-e and Supplementary Fig. 3). In addition, the magnitude of I_NMDA_ measured in the absence and presence of PEAQX and AV-5 did not differ between WT and Pah^Enu2^ mice (Fig. 2f,g). Furthermore, four episodes of theta-burst stimulation (4X TBS) of the SC axons induced similar magnitudes of LTP in both genotypes (Fig. 2h). Surprisingly, however, perfusion of L-Phe (250 μM) during the TBS significantly attenuated LTP induction (Fig. 2h), despite that L-Phe had no effect on the fiber volley amplitudes, field excitatory postsynaptic potential (fEPSP) slopes, and population spikes in the basal condition (Supplementary Fig. 4). Moreover, the extent of LTP attenuation by L-Phe perfusion did not differ between the WT and Pah^Enu2^ slices. Consistent with I_NMDA_ facilitation at 10 μM glycine (Fig. 1j), L-Phe-induced attenuation of LTP was observed under treatment with 10 μM glycine in the ACSF (Supplementary Fig. 5a,b). These results indicate that Phe in the CSF has a greater impact on LTP impairment than altered brain development^36,37^ caused by chronic exposure to elevated Phe in Pah^Enu2^ mice.

**Fig. 2:**
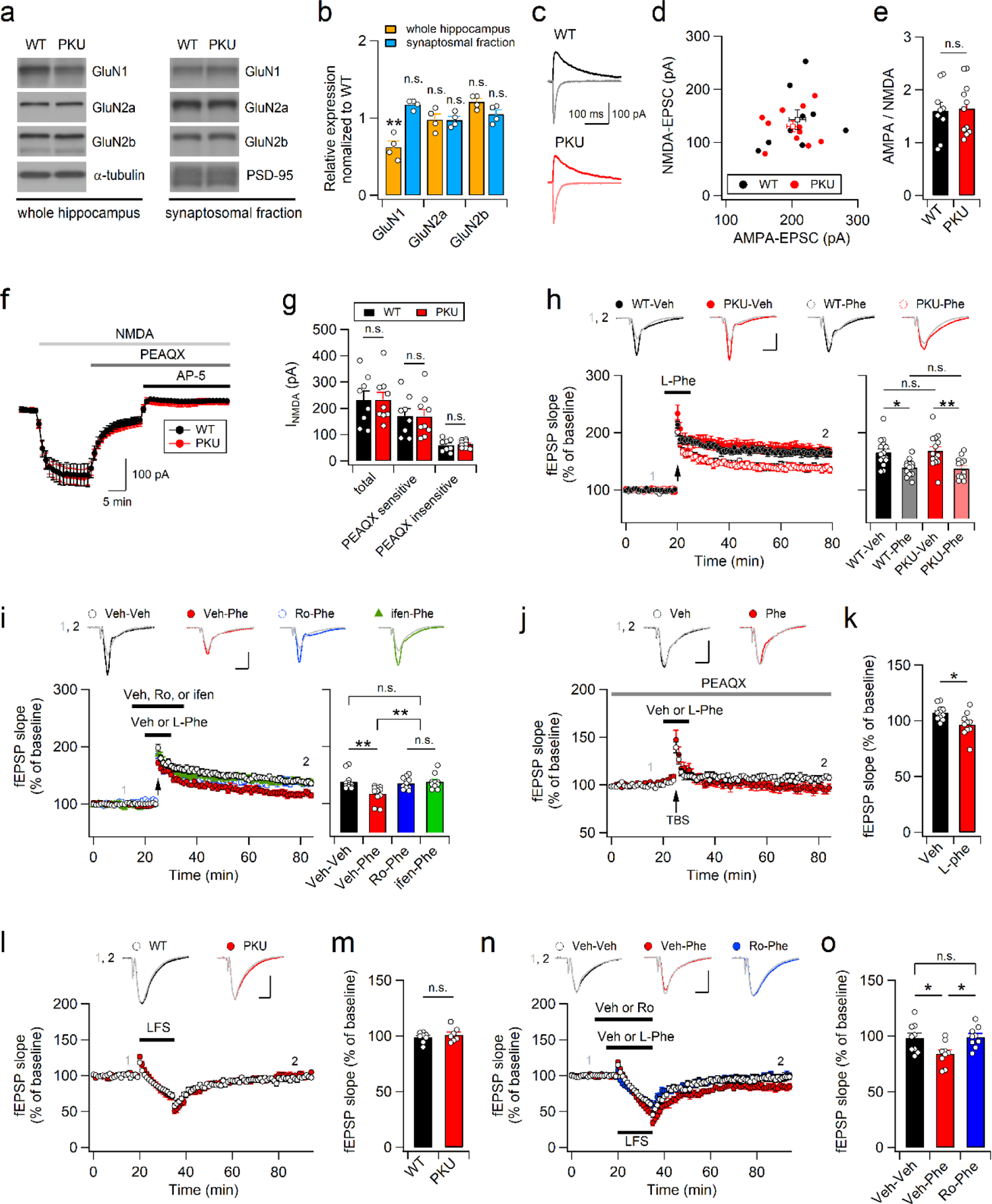
L-Phe perturbs synaptic plasticity though the activity-dependent down-regulation of AMPARs. **a, b**, Representative western blots (**a**) and expression levels (**b**) of NMDAR subunits in the total and synaptosomal fractions of the WT and Pah^Enu2^ hippocampi. **c**, Representative AMPAR- and NMDAR-EPSCs measured at −70 and +40 mV, respectively, in CA1 pyramidal neurons. **d**, The peak amplitude of NMDAR-EPSCs were plotted against AMPAR-EPSCs. **e**, Normal AMPA-NMDA ratios in the CA1 pyramidal cells of Pah^Enu2^ mice. **f**, PEAQX (0.5 μM) reduced I_NMDA_ and subsequent AP-5 (50 μM) perfusion blocked the remnant. **g**, Total I_NMDA_ and PEAQX-sensitive and -insensitive components after sequential application of PEAQX and AP-5 in CA1 pyramidal cells of WT and Pah^Enu2^ mice. **h**, L-Phe reduces the magnitude of LTP in both WT and Pah^Enu2^ mice. (top) Representative traces of fEPSP obtained at the indicated time points. (bottom, left) fEPSP slopes were normalized to those obtained in the baseline and plotted against time. L-Phe was perfused from 5 min before to 1 min after TBS (arrow). (bottom, right) The magnitudes of LTP measured in the absence and presence L-Phe in each genotype. **i**, GluN2B antagonists block the effect of L-Phe on the TBS (arrow)-induced LTP. Sample traces (top), time course of fEPSP slopes (bottom, left), and the magnitude of LTP (bottom, right) in each condition. **j**, PEAQX blocks LTP induction, and L-Phe perfusion during the peri-TBS period induces an LTD-like decrease in fEPSP slopes. Top, sample traces of fEPSPs. **k**, fEPSP slopes during the last 10 min were normalized to baseline. **l**, **m**, WT and Pah^Enu2^ slices exhibit similar responses to LFS. **m**, Normalized fEPSP slopes during the last 10 min in WT and Pah^Enu2^ slices. **n**, Ro blocks the effect of L-Phe on LTD facilitation. Sample traces of fEPSPs (top). **o**, Normalized fEPSP slopes during the last 10 min in each condition. (**h**-**j**, **l**, and **n**) Scale bars, 5 ms and 0.5 mV.

We hypothesized that CSF Phe hyperactivates extrasynaptic GluN2B-containing NMDARs during TBS, consequently inducing homeostatic downregulation of AMPARs^38^. Indeed, treatment of slices with Ro (2 μM) or ifen (6 μM) completely blocked the effect of L-Phe on LTP attenuation in the WT slices (Fig. 2i). Meanwhile, PEAQX blocked LTP induction such that the increased fEPSPs caused by TBS rapidly returned to baseline levels. Under these conditions, perfusion of L-Phe during TBS induced an LTD-like decrease in synaptic strength, indicating the downregulation of AMPARs (Fig. 2j,k). Consistent with this, neither LTP nor L-phe-induced synaptic depression was observed after the TBS when the slices were co-treated with PEAQX and Ro (Supplementary Fig. 5c,d).

We further determined the synaptic responses of Pah^Enu2^ hippocampus to low-frequency stimulation (LFS), which is widely used to induce LTD in juvenile mice. Although LFS (1 Hz, 900 stimulations) did not induce LTD in either genotype, the fEPSP slopes in WT and Pah^Enu2^ mice decreased to the same extent during LFS and gradually returned to baseline levels (Fig. 2l,m). Notably, L-Phe perfusion promoted synaptic depression during LFS and induced stable LTD, which was blocked by Ro perfusion during LFS (Fig. 2n,o). Collectively, these results suggest that the concentration of Phe observed in PKU CSF hyperactivates GluN2B-containing receptors and induces activity-dependent downregulation of AMPARs.

### L-Phe challenge recapitulates cognitive symptoms of PKU in adult mice

Pah^Enu2^ mice exhibit different behavioral phenotypes depending on their genetic background, despite similar biochemical phenotypes^39–41^. Consistent with the results of previous studies^39,42^, we observed impaired learning and memory in Pah^Enu2^ mice with a C57BL/6N background. Pah^Enu2^ mice behaved normally in the open-field test (OFT), but displayed impaired performance in novel object recognition (NOR), object location memory (OLM), and Morris watermaze (MWM) tests (Supplementary Fig. 6).

If Phe_CSF_ exerted a profound effect on cognitive dysfunction in Pah^Enu2^ mice, L-Phe challenge would induce similar phenotypes in adult WT mice. Intraperitoneal (i.p.) administration of L-Phe at 1 mg/g of body weight reported to increase the Phe_Serum_ to 1.7 ± 0.1 mM in WT mice^43^, and i.p. injections to rats with the similar concentration (10 mmole/kg) induced, within 1 h, the Phe_Brain_ comparable to those observed in patients with PKU at autopsy^44^. As the phosphorylation status of eukaryotic elongation factor 2 (eEF2) changes rapidly in response to neuronal activity^45^, we examined the effect of L-Phe challenge on the proportion of phosphorylated eEF2 (p-eEF2) in total eEF2. L-Phe (1 mg/g)-treated mice exhibited an increased p-eEF2/eEF2 ratio in both the hippocampus and whole brain compared to vehicle treated mice (Fig. 3a). Enhanced eEF2 phosphorylation was also detected in the Pah^Enu2^ hippocampus (Fig. 3b), indicating that both Pah^Enu2^ and L-Phe-treated mice exhibited reduced neuronal activity in the brain. To directly monitor changes in neuronal activity induced by L-Phe challenge, we performed fiber photometry recordings in freely behaving mice, and observed a reduction in neuronal activity in the medial prefrontal cortex (mPFC) following L-Phe administration (Fig. 3c-e). These results indicate that L-Phe challenge modifies neuronal activity in WT mice similar to those of the Pah^Enu2^ mice. Intriguingly, pretreatment (30 min before L-Phe injection) with Ro abolished the effect of L-Phe on hippocampal eEF2 phosphorylation (Fig. 3f), indicating an association between GluN2B receptor activation and L-Phe-induced reduction of neuronal activity. In support of this idea, fiber photometry detected a reduction in and recovery of neuronal activity in the mPFC following L-Phe and subsequent Ro injections (Fig. 3g,h).

**Fig. 3:**
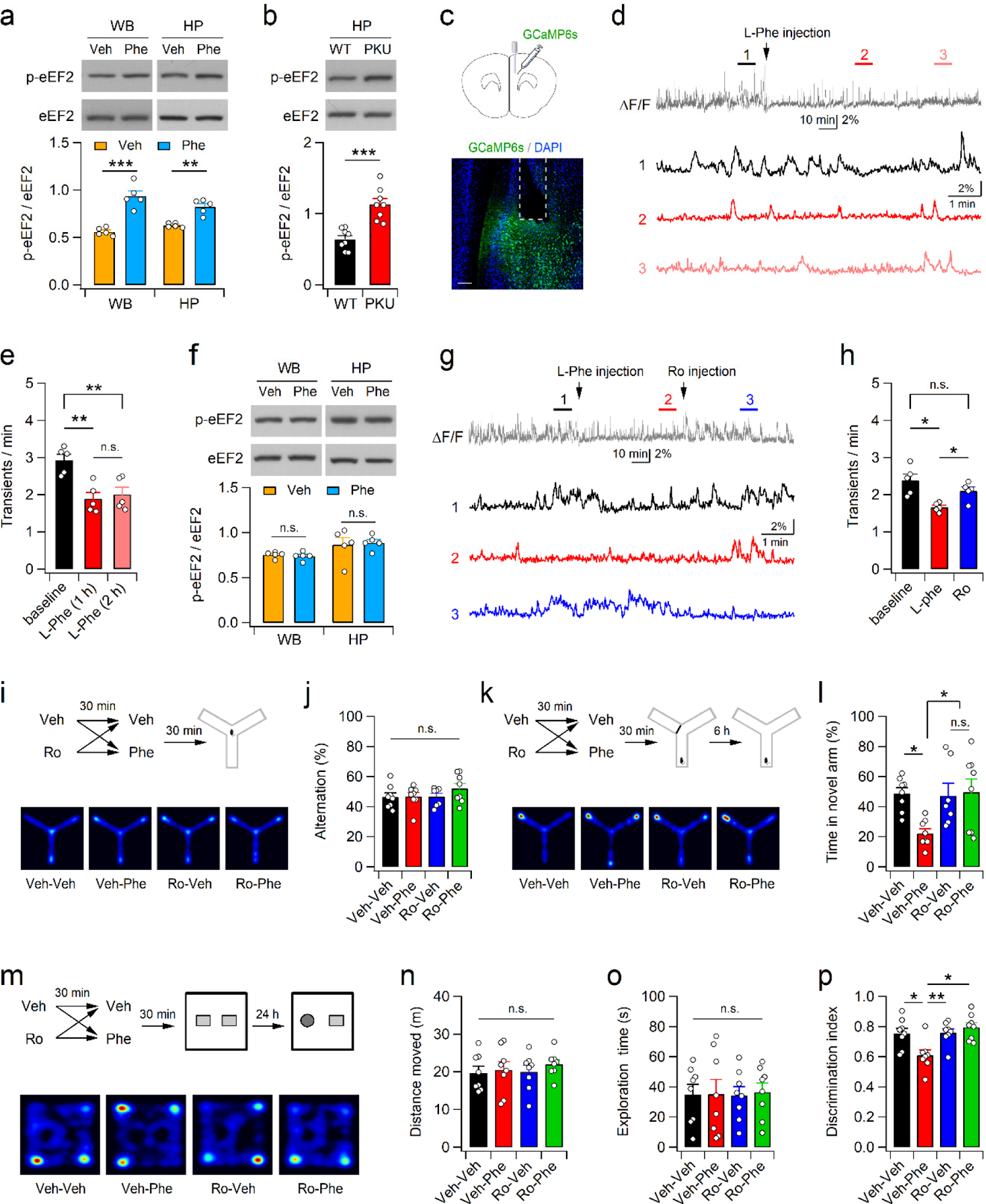
L-Phe loading decreases neural activity and impairs learning. **a**, Western blot analyses for the protein levels of p-eEF2 and EF2 were performed 30 min after vehicle (veh) or L-Phe (1 mg/g, i.p.) administration. WB, whole brain, HP, hippocampus. (bottom) Quantification of the p-eEF2/eEF2 ratio. **b**, Enhanced eEF2 phosphorylation (top) and the p-eEF2/eEF2 ratio (bottom) in the Pah^Enu2^ hippocampus. **c**, Experimental design for fiber photometry recoding in the mPFC of WT mice. Bottom, immunohistochemical staining of a mPFC section showing the GCaMP6s-expressing cells (green) and the canula placement. DAPI (blue) was used to identify the brain structures. Calibration, 200 μm. **d**, Neuronal activity of CaMKII-expressing cells in the mPFC was decreased by L-Phe administration. Bottom shows the fluorescence signals obtained during the indicated periods (1, 2, 3) on expanded time scale. **e**, Quantification of the frequency of Ca^2+^ transients obtained from 5 mice. **f**, Ro blocks L-Phe-induced eEF2 phosphorylation in the hippocampus. Ro (3 mg/kg, i.p.) was administered 30 min before L-Phe or vehecle injection. **g**, L-Phe and Ro were sequentially administered during the recording. **h**, Summary of the frequency of fluorescence transients during the baseline and perfusion of L-Phe and Ro. **i**, Experimental design (top) and activity paths (bottom) of mice in the Y-maze. **j**, The percentage of spontaneous alternation in the Y-maze measured 30 min after L-Phe or vehicle injections. **k**, Mice received L-Phe or vehicle 30 min before training (top). Bottom, activity paths of mice during the test session. The test session was conducted 6 h after the training session. **l**, Mice treated with L-Phe spent significantly less in the novel arm. **m**, Ro and L-Phe were administered 1 h and 30 min before the training session of NOR, respectively. (bottom) Activity paths of mice during the test session of NOR are shown by heat maps. **n**, **o**, Quantification of distance moved (**n**) and time spent exploring a novel object or a familiar object (**o**) for 10 min during the test session of NOR. **p**, Relative preference for the novel object was calculated using a discrimination index.

L-Phe challenge did not affect spontaneous alteration of WT mice during the Y-maze test, indicating normal working memory (Fig. 3i,j). However, L-Phe treatment 30 min before the training phase of the novel arm exploration significantly impaired novel arm discrimination of mice during the test phase that performed 6 h after the training phase (Fig. 3k,l). Similarly, mice received L-Phe 30 min before the sample phase of the NOR test spent less time exploring the novel object than vehicle treated mice in the test session, which was performed 24 h after the sample phase (Fig. 3m-p). Considering that serum L-Phe concentrations return to normal within 2 h of administration^43^, L-Phe might have influenced learning more than recall performance of mice. Importantly, pretreatment (30 min before L-Phe administration) of mice with Ro (3 mg/kg) abolished the L-Phe-induced impairment in novel arm discrimination and NOR (Fig. 3k-p). We confirmed that the doses of L-Phe and Ro used in this study did not affect general activity or explorative behaviors of mice in the OFT (Supplementary Fig. 7). Collectively, these results suggest that L-Phe elevation is sufficient to impair learning and memory, and that GluN2B receptors play a key role in L-Phe-induced cognitive dysfunction.

### Suppression of GluN2B rescues impaired learning in Pah^Enu2^ mice

Since excitatory synaptic transmission and plasticity in the Pah^Enu2^ hippocampal slices did not differ from those of WT slices in normal ACSF, we wondered whether suppression of GluN2B would restore impaired learning and memory in adult Pah^Enu2^ mice. We examined the effects of GluN2B antagonists on the NOR of Pah^Enu2^ mice and found that the administration of ifen (5 mg/kg, i.p.) or Ro (3 mg/kg, i.p.) 30 min before the sample session significantly improved the performance of NOR in Pah^Enu2^ mice during the test session (Fig. 4a-f). Consistent with this observation, the administration of Ro increased the activity of CaMKII-expressing neurons in the Pah^Enu2^ mPFC, as detected by fiber photometry (Supplementary Fig. 8). The administration of ifen also improved OLM in PKU mice (Fig. 4g-i), further confirming the effect of GluN2B suppression on learning performance in PKU mice. Ifen did not induce abnormal behavior in WT or Pah^Enu2^ mice during the OFT (Supplementary Fig. 9).

**Fig. 4:**
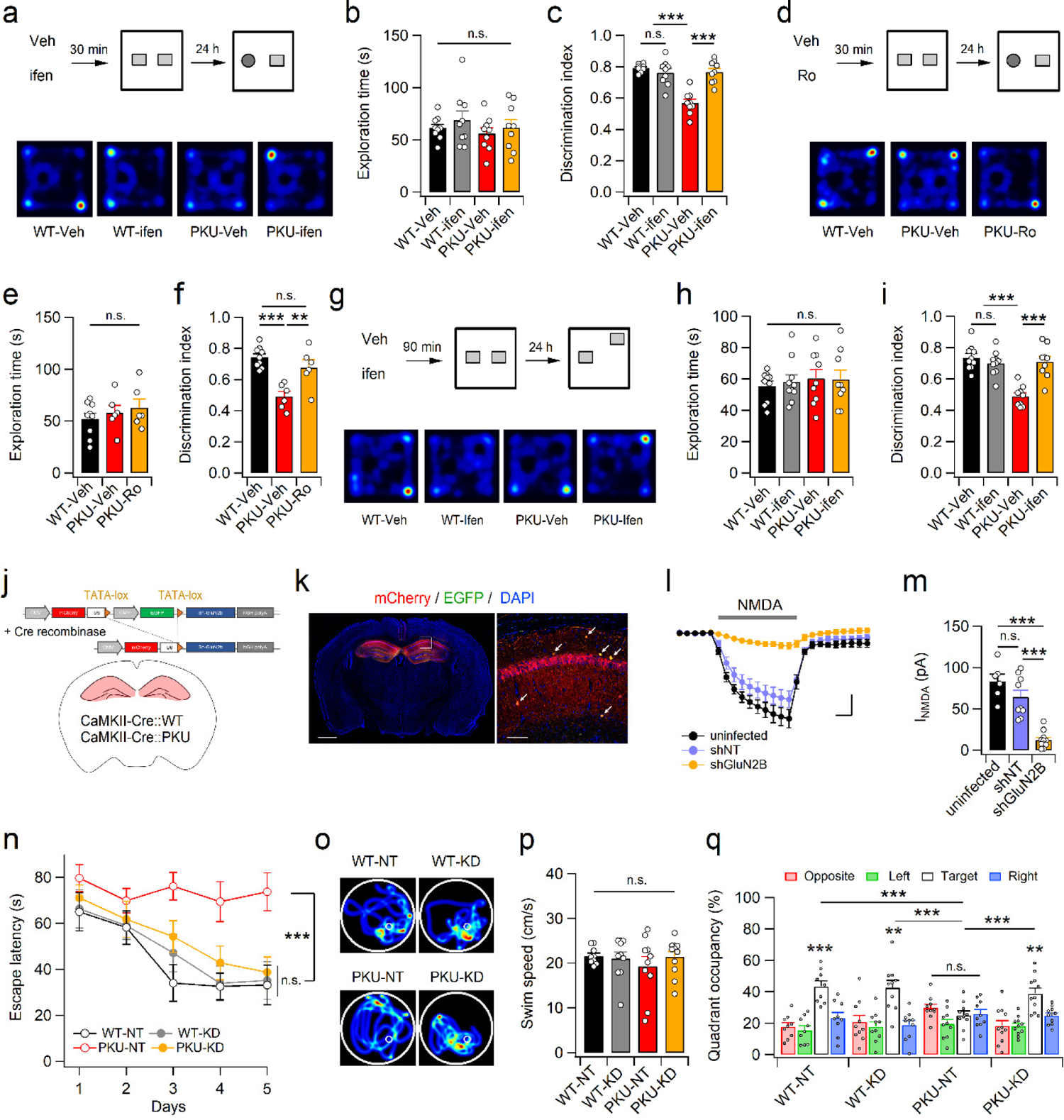
GluN2B suppression improves learning and memory impairment in Pah^Enu2^ mice. **a**, Mice were treated with Veh or ifen 30 min before the NOR training session. Bottom, example path recordings of mice during the NOR test session. **b**, **c**, The time spent exploring the two objects (**b**) and preference for novel object (**c**) are summarized. **d**-**f**, Same as **a**-**c**, but for Ro. **g**, Experimental design (top) of the OLM test and sample path recordings (bottom) of mice during the OLM test session. **h**, **i**, Quantification of time spent exploring the two objects (**h**) and preference for the moved object (**i**) during the OLM test session. **j**, Design of Cre-dependent expression of shGluN2B in CaMKII-expressing cell in the dorsal hippocampus. **k**, Immunohistochemical staining of hippocampal section shows expression of mCherry (red) in the hippocampus. Calibration, 1 mm. Right, magnified image of the region corresponding to the white box in the left panel showing the absence of an EGFP signal in CA1 principal cells. Yellow cells (arrows) indicate putative interneurons expressing both EGFP and mCherry. Calibration, 100 μm. **l**, I_NMDA_ in the CA1 pyramidal neurons was measured in the presence of blockers for GluN2A, Na^+^-channels, AMPARs, and GABA_A_Rs. Scale bars, 2 min and 20 pA. **m**, Reduced I_NMDA_ in CA1 pyramidal neurons expressing shGluN2B. **n**, Escape latency of mice in each group during the 5-day training session of MWM test. **o**-**q**, Representative swim path (**o**), swim speed (**p**), and quadrant occupancy (**q**) of mice during the MWM probe trials.

We wondered whether Glu2B suppression could rescue the impaired MWM performance of Pah^Enu2^ mice. To stably suppress the function of GluN2B receptors, we generated an adeno-associated virus (AAV) carrying a Cre-dependent short hairpin RNA (shRNA) expression vector that targets GluN2B (Fig. 4j). Infection of the shGluN2B virus into the dorsal hippocampus of CaMKII-Cre mice (Fig. 5k) induced a significant reduction in the expression levels of GluN2B but not GluN1, GluN2A, or PSD-95 in the synaptosomal fraction (Supplementary Fig. 10 and 11). We further confirmed a significant reduction in the GluN2B-mediated current in CA1 pyramidal neurons infected with the virus expressing shGluN2B compared to those infected with the non-targeting scrambled sequence (shNT) or uninfected (Fig. 4l,m).

**Fig. 5:**
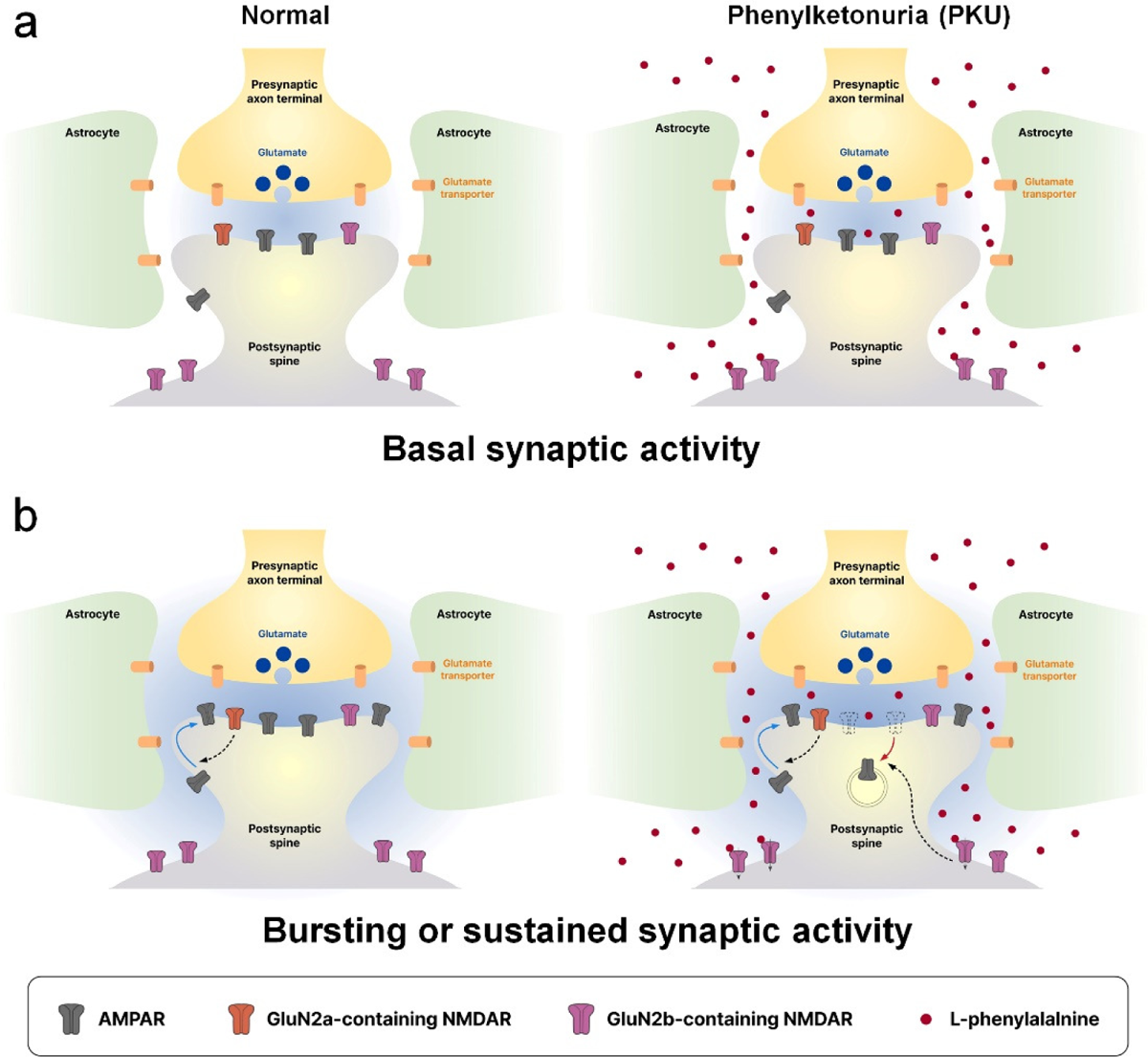
A model of synaptic disturbance in PKU. **a**, Phe preferentially affects glutamate-induced activation of GluN2B-containing NMDARs. During the low frequency activity, extrasynaptic NMDARs are minimally affected by Phe, because the activity of glutamate transporters limits the diffusion of glutamate onto extrasynaptic NMDARs. **b**, Bursting or sustained synaptic activity induces glutamate spillover and activation of extrasynaptic NMDARs. Phe hyperactivates GluN2B-containing NMDARs, consequently inducing activity-dependent downregulation of AMPARs.

Infection of the dorsal hippocampus with shGluN2B did not significantly affect MWM performance in adult CaMKII-Cre mice (WT-KD). Meanwhile, CaMKII-Cre;Pah^Enu2^ mice infected with shGluN2B (PKU-KD) exhibited significantly enhanced learning during the 5-day training period, whereas CaMKII-Cre;Pah^Enu2^ mice expressing scrambled sequences (PKU-NT) did not show any signs of learning during the same period (Fig. 4n). Consistent with this finding, CaMKII-Cre;Pah^Enu2^ mice expressing shGluN2B exhibited quadrant occupancy similar to that of CaMKII-Cre mice during the probe trial (Fig. 4o-q). Collectively, these results suggest that GluN2B suppression improves cognitive function in adult PKU mice.

## Discussion

In this study, we investigated the synaptic mechanisms underlying cognitive impairment associated with PKU. We provided evidence that Phe predominantly affects GluN2B-rather than GluN2A-containing NMDARs. Importantly, we found that the concentration of Phe in the CSF of patients with PKU, in contrast to that observed in the serum, upregulated the activity of GluN2B receptors. Based on its diffusion coefficient^46^, Phe transported across the BBB is thought to be distributed evenly in both the synaptic and extrasynaptic areas of the brain. GluN2B receptors are located predominantly, but not exclusively, in the extrasynaptic area at the mature synapse^28,47^, and the activity of glutamate transporters limits the activation of extrasynaptic NMDARs during low-frequency activity^48^. However, bursting or sustained synaptic activity results in glutamate spillover, which activates extrasynaptic NMDARs^49^. Phe present in PKU CSF may induce hyperactivation of GluN2B-containing NMDARs, and consequently lead to abnormal temporal integration of synaptic inputs and activity-dependent downregulation of AMPARs^38,47^ (Fig. 5).

Based on the finding that L-Phe at concentrations observed in or higher than PKU serum reduced I_NMDA_ in cultured hippocampal neurons^17,18^, it is widely thought that elevated Phe in PKU inhibits NMDAR function. While the Phe concentration is higher than 1 mM in the serum, clinical studies have shown that the Phe concentration in the PKU CSF is much lower than that in serum, as Phe crosses the BBB through L-type amino acid transporter 1 (LAT1)-mediated transport^2^. Blood Phe concentrations (Phe_Blood_) of 1.0 mmol/L were consistently associated with brain Phe concentrations (Phe_brain_) of 0.2 – 0.3 mmol/kg in humans^21^, and the linear regression model predicts the relationship between Phe_brain_ and Phe_Blood_ as follow (in μM): Phe_Brain_ = 22.02 + 0.22 × Phe_Blood_^24^. A recent preclinical study also reported that PKU (Pah^enu2^) mice exhibit a strong correlation between Phe_brain_ and Phe_Plasma_ and that Phe_brain_ in the C56BL/6 and BTBR strains of PKU mice were 0.298 and 0.392 mmol/kg, respectively^25^.

Although the pathogenesis of the neuropsychological complications in PKU remains unclear, abnormal brain development under elevated Phe concentrations is believed to be associated with intellectual disability, seizures, microcephaly, and psychiatric symptoms in patients with PKU^50,51^. A reduction in myelination, dendritic arborization, and synaptic spines has been observed in patients with PKU^9^. However, the myelination status^52,53^ and spine density^36,37,54^ in Pah^enu2^ mice remain controversial, and opposing results has been reported. Moreover, in vitro studies have found that incubation of mouse cerebellar organotypic slices with 1.2 and 2.4 mM, but not 0.6 mM, Phe reduced myelination^55^, and that 1 and 2 mM Phe increased dendritic arborization^56^ and spine density^37^, respectively, in hippocampal neurons.

Synaptic NMDARs in the central nervous system undergo replacement of their subunit composition, predominantly from GluN2B to GluN2A, during early postnatal development^47,57^. At immature glutamatergic synapses, GluN2B contributes to synaptic NMDAR signaling, and the activity of GluN2B negatively regulates AMPAR incorporation^58,59^. Hence, in contrast to the adult brain, elevated Phe levels may enhance the electrical and biochemical signaling of synaptic NMDARs in the neonatal brain. Hyperactivation of synaptic GluN2B may delay the development of glutamatergic synapses and brain maturation by hampering the incorporation of AMPARs into immature synapses. Untreated PKU results in severe intellectual impairment, and late treatment with a low-Phe diet partially reverses this cognitive impairment^1,60^, indicating that the neonatal brain, in which GluN2B is predominant^57^, is more sensitive to elevated Phe levels than the mature brain. Importantly, a recent study reported that the overexpression of GluN2B reduced both tonic GABA currents and the surface expression of α5 subunits of GABA_A_Rs (α5-GABA_A_R) in hippocampal neurons^61^. Although the expression levels of α5-GABA_A_R and epileptic mechanisms are unknown in patients with PKU, epilepsy is frequently accompanied by PKU. Adult (18-20 weeks) but not younger (5-7 weeks) Pah^Enu2^ mice of the BTBR strain also exhibited enhanced susceptibility to audiogenic seizures compared to WT mice^62^. However, in our study, spontaneous seizure behavior was not observed in Pah^Enu2^ mice.

One of the important findings of this study was that Phe at the concentration observed in PKU CSF perturbs synaptic plasticity through the hyperactivation of GluN2B-containing NMDARs. Similar to our results, amyloid-β protein (Aβ) has been suggested to hyperactivate extrasynaptic GluN2B receptors and consequently facilitate LTD but impair LTP in the hippocampus^63,64^. L-Phe is unlikely to inhibit glutamate reuptake or promote glutamate spillover, but acts directly on GluN2B receptors, as L-Phe did not affect NMDAR-EPSCs in the absence of TBOA, and L-Phe increased I_NMDA_ in HEK293 cells expressing GluN1.1a and GluN2B. In addition to possible structural changes in the brain^15^, the hyperactivation of GluN2B and/or perturbed synaptic plasticity may contribute to cognitive deficits in adults with PKU. In support of this idea, the L-Phe challenge in adult WT mice impaired the NOR and OLM performance, which was abrogated by pretreatment with GluN2B inhibitors. A recent clinical study reported that early treated adults with PKU demonstrated underperformance in cognitive tests, including processing speed, executive function, and learning, and that processing speed was significantly related to Phe concentration at the time of testing^3^. In addition, early treated adults with lower Phe levels performed better than those with higher Phe levels in most cognitive domains including IQ^4^. These observations indicate that elevated Phe levels still influence cognitive function in adults^15^, and that dietary control alone may not be sufficient to prevent suboptimal cognitive outcomes^3^. In fact, a low-Phe diet is difficult to maintain, and compliance in adolescent and adults is often poor. Tetrahydrobiopterin and large neutral amino acid (LNAA) treatments are known to have beneficial effects on cognitive function^1^. However, there are substantial unmet needs for patients with PKU. Notably, CSF analyses revealed a significant enhancement in the levels of Aβ_1-42_, total tau, and phosphorylated tau in early-treated patients with PKU compared to healthy controls^15^. These observations further support the therapeutic potential of GluN2B blockers in the treatment of neurological complications of PKU. The reversal of impaired learning and memory in adult Pah^Enu2^ mice by both pharmacological treatment and shRNA-mediated gene silencing further indicates that cognitive impairment in adults with PKU is treatable and preventable by restoring GluN2B activity and synaptic plasticity. Further investigations are needed to determine the association between GluN2B signaling and neuropsychological complications in individuals with PKU.

## Methods

### Animals

All experiments were performed using C57BL/6N mice of both sexes (Orient Bio, Sungnam, Korea), except for L-Phe challenge, which were performed exclusively with male mice. Animals were group-housed (3−5/cage) in a specific pathogen-free facility and maintained in a climate-controlled room under a 12 h light/dark cycle. All mice had free access to water and standard chow (20% protein containing 0.98% L-Phe). Pah^Enu2^ (Jackson Laboratory stock #002232) mice were kindly provided by Sung-Chul Jung (Ewha Womans University) and backcrossed with C57BL/6N mice for at least 5 generations before use. The genotypes of Pah^Enu2^ mice were determined by restriction endonuclease digestion of the PCR product. Genomic DNA was extracted from mouse tail, and exon 7 of the Pah gene was amplified using oligonucleotide primers 5′-CCTTGGGGAGTCATACCTCA-3′ and 5′-ATAAAGCAGGCAGTGGATCA-3′. The 317-bp PCR product was digested with the restriction endonuclease MboII overnight at 37 °C, and the restriction fragments were separated by electrophoresis on a 2% acrylamide gel. CaMKII-Cre transgenic mice were kindly provided by Yong-Seok Lee (Seoul National University), and were backcrossed with C57BL/6N mice for at least 10 generations before use. All animal maintenance and experiments were approved by the Institutional Animal Care and Use Committee (IACUC) of SNU.

### Electrophysiology

Parasagittal hippocampal slices (400 µm thick) were prepared using a vibratome (Leica, Germany) in ice-cold dissection buffer (sucrose 230 mM; NaHCO_3_ 25 mM; KCl 2.5 mM; NaH_2_PO_4_ 1.25 mM; D-glucose 10 mM; Na-ascorbate 1.3 mM; MgCl_2_ 3 mM; CaCl_2_ 0.5 mM, pH 7.4 with 95% O_2_/5% CO_2_). Immediately after sectioning, the CA3 region was surgically removed. The slices were allowed to recover at 36°C for 1 h in normal ACSF (NaCl 125 mM; NaHCO_3_ 25 mM; KCl 2.5 mM; NaH_2_PO_4_ 1.25 mM; D-glucose 10 mM; MgCl_2_ 1.3 mM; CaCl_2_ 2.5 mM, pH 7.4 with 95% O_2_/5% CO_2_), and then maintained at room temperature.

Slices were placed in a submerged recording chamber, which was perfused continuously with heated (29–30°C) ACSF. All electrophysiological recordings were performed using a MultiClamp 700B amplifier and Digidata 1440A interface (Molecular Devices, San Jose, CA, USA). The signals were filtered at 2.8 kHz and digitized at 10 kHz. The data were analyzed using custom macros written in Igor Pro (WaveMetrics).

To measure I_NMDA_ in CA1 neurons at a holding potential of –40 mV, whole-cell voltage clamp recordings were made using patch pipettes (3–4 MΩ) filled with solution containing (in mM) 100 Cs-gluconate, 10 TEA-Cl, 10 CsCl, 8 NaCl, 10 HEPES, 4 Mg-ATP, 0.3 Na-GTP, 0.5 QX-314-Cl and 10 EGTA, adjusted to pH 7.25 and 290 mOsm/kg. Picrotoxin (50 µM), NBQX (10 µM), and TTX (1 µ M) were added to the ACSF. NMDA-EPSCs were measured using the same pipette solution used for measurement of I_NMDA_. Picrotoxin (50 µM) and NBQX (10 µM) were added to the ACSF. Synaptic responses were evoked at 0.05 Hz with an ACSF-filled broken glass pipette (0.3–0.5 MΩ) placed in the proximal region of the stratum radiatum. The series and seal resistances were continuously monitored using short (50 ms) test (2 mV) pulses, and data were discarded if they changed by more than 20% during the recordings.

To measure the synaptic AMPA/NMDA ratio, AMPAR-mediated EPSCs were obtained by averaging 30–40 traces recorded at –70 mV. The stimulation intensity was adjusted to yield a 100–300 pA EPSC peak amplitude. After recording AMPAR-mediated EPSCs, NBQX (10 µM) was added to ACSF, and 30–40 traces of NMDAR-mediated EPSCs were recorded at +40 mV.

To record field excitatory postsynaptic potential (fEPSP) at SC-CA1 synapse, a recording (3−4 MΩ) pipette filled with ACSF was placed in the stratum radiatum. Synaptic responses were evoked by stimulating Schaffer collaterals (SCs) with an ACSF-filled broken glass pipette (0.05 Hz), and the stimulation intensity was adjusted to yield approximately 30% of maximal responses. LTP was elicited by four trains of theta-burst stimulation (TBS), with 10 s intertrain interval. The TBS consisted of 10 bursts, each consisting of 4 pulses at 100 Hz, with an interburst interval of 200 ms. The low-frequency stimulation consisted of 900 pulses at 1 Hz. Slices displaying unstable (10%) baseline recordings were excluded from the analysis. Extracellular population spike recordings were performed by using a recording electrode placed on the CA1 stratum pyramidale. Synaptic responses were evoked by a stimulating electrode placed in the stratum radiatum.

### Cell culture and transfection

The human embryonic kidney cell line (HEK-293T) cells was cultured on poly D-lysine-coated glass coverslips in high-glucose (4.5 g/L) Dulbecco’s modified Eagle’s medium (DMEM) containing 4 mM L-glutamine, 3.7 g/L sodium bicarbonate, 10% (v/v) fetal bovine serum (FBS), 1 mM sodium pyruvate, and 1% (v/v) penicillin/streptomycin. After transfection, the cell culture medium was replaced with fresh medium containing D-AP5 (50 µM) and 1 mM MgCl_2_. hGluN1.1a and hGluN2b had been previously described^65^. Plasmids were transfected at the DNA ratio of hGluN1.1a, hGluN2b, and EGFP with 2:2:1 (total 0.5 µg / coverslip) using the Lipofectamine 3000 transfection reagent kit (Thermo Fisher Scientific). Transfected cells were identified by EGFP signals, and electrophysiological recordings were performed 24−72 h after transfection.

### Drugs

L-Phe was purchased from Sigma-Aldrich (St. Louis, MO, USA). To prepare stock solution, L-Phe was dissolved to 1 M in 100 ml of distilled water using NaOH at 37°C, and the stock solution (pH 7.4) was aliquoted and cryopreserved until use. All reagents were purchased from Sigma-Aldrich (St. Louis, MO, USA), except for PEAQX, Ro25-6981, ifenprodil, QX-314-Cl, NBQX, TBOA, and AP-5, which were purchased from Hello Bio (Bristol, UK).

### Behavioral analyses

Behavioral tests were performed between 10 a.m. and 6 p.m. on mice that were at least 8 weeks old. All experimental mice were acclimated to the behavior testing room for at least 1 h prior to testing. The testing apparatus was cleaned with 70% ethanol between trials.

The open field test (OFT) was conducted using open field boxes (40 × 40 × 40 cm) with opaque walls in a dimly lit room. The mice were placed in the center of an open field box, and the behavior of each mouse was monitored using video recordings. The total distance traveled in the entire box and the time spent in the center zone (20 × 20 cm) were calculated using video tracking software (Ethovision XT, Noldus, Netherlands).

The NOR and OLM tests were conducted in the same box used for the OFT, but with two identical objects in the middle of the box. During the acquisition (sample) session of the NOR and OLM tests, each mouse was placed in the center of the box and allowed to explore the two objects for 10 min. The NOR and OLM test sessions were conducted 24 h after the acquisition session. During the NOR test session, mice were returned to a box in which one of the familiar objects was replaced with a new object. To minimize any bias in the location of objects, the relative locations of familiar and novel objects were counterbalanced between trials. The behavior of each mouse was monitored for 10 min using video recordings, and object interaction was defined as sniffing, brief contact, and/or approaching an object. The discrimination index (%) of the NOR test was calculated as follows: [(novel object interaction)/(familiar object interaction + novel object interaction)] × 100. During the OLM test session (10 min), mice were returned to the arena, where one of the two familiar objects was moved to the corner of the box. The position of the moved object was counterbalanced between the mice. The discrimination index (%) of the OLM test was calculated as follows: [(moved object interaction) / (unmoved object interaction + moved object interaction)] × 100.

Y-maze tests were conducted using a symmetrical, top-open, Y-shaped maze with acrylic walls. Each arm of the Y-maze was 35 cm long × 5 cm wide × 13 cm tall. The mice were allowed to explore all three arms for 30 min, and spontaneous alternations in each mouse were analyzed from the initial 10 min of the habituation period. Spontaneous alternations (%) were defined as consecutive entries into three different arms (e.g., ABC or BAC, but not ACA) of the Y-maze divided by the number of possible alternations: [number of alternations / (number of total arm entries – 2) ×100]. During the training session of the spatial reference memory test, one of the three arms was blocked using an acrylic baffle. Each mouse was placed in the starting arm and allowed to explore both the starting and other arms freely for 15 min. The test session of the spatial reference memory test was conducted 6 h after the training session. The acrylic baffle was removed, and mice were allowed to explore all three arms of the Y-maze for 5 min. The percentage of time spent in each arm was analyzed using a video tracking software (Ethovision XT).

The MWM test was performed using a white circular pool (120 cm in diameter) filled with warm (24°C−25°C), opacified water. The pre-training session (2 days) consisted of handling for 5 min and acclimation on a visible platform (10 cm in diameter) for 2 min, with 1 trial per day. During the 5 days-training period, mice were allowed to find the hidden platform with 3 consecutive trials per day and were guided to the platform if they failed to find the platform within 90 s. The starting position (opposite quadrant, right adjacent quadrant, left adjacent quadrant) of each mouse was alternated between trials in a pseudo-random order. Each mouse was allowed to remain on the platform for 30 s, followed by a 30-s rest in its home cage. The mice were allowed to find the hidden platform from a different starting point. The probe test was performed 24 h after the completion of the training trials, and the mice were allowed to swim for 60 s in the absence of a hidden platform. The escape latency, swim distance, speed, and swim pattern were analyzed using video tracking software (Ethovision XT).

### Surgery and stereotaxic injection

Mice were deeply anesthetized with a mixture of Zoletil (50 mg/kg, i.p.) and xylazine (1 mg/kg, i.p.) and placed in a stereotaxic device. The skin was cut over the midline and craniotomies were performed bilaterally over the dorsal hippocampus (−1.9 anteroposterior, ±1.5 mediolateral, −1.5 dorsoventral from the bregma and dura). AAV vectors expressing GluN2B shRNA (5′-TgtaccaacaggtctcaccttaaacTTCAAGAGAgtttaaggtgagacctgttggtacTTTTTTC-3′) and scrambled shRNA (5′-TaccatcttgacataagcgacctcaTTCAAGAGAtgaggtcgcttatgtcaagatggtTTTTTTC-3′) were kindly provided by Ronald Duman (Yale School of Medicine), and AAVs were produced and titrated by the Stanford University Gene Vector and Virus Core.

Purified AAV (0.5 µ L/side) was injected using a Hamilton syringe at a rate of 100 nL/min. After completion of the injection, the needle (33 gauge) was maintained in place for an additional 10 min to allow diffusion of the injection medium before being carefully retracted to prevent backflow. Experiments were performed 3-4 weeks after the viral injections.

### Fiber photometry

Under deep anesthesia with a mixture of Zoletil and xylazine, AAV5/Syn-Flex-GCaMP6s-WPRE-SV40 (Addgene, Cat. #100845) were unilaterally injected into the mPFC (+1.8 anteroposterior, ±0.5 mediolateral, −2.5 dorsoventral), after which a cannula (400 μm diameter, 0.39 NA, CFM14U-20, Thorlabs) for fiber photometric recording was implanted above the injection site. The implants were then secured using dental cement and metal screws anchored to the skull of the contralateral hemisphere. The mice were allowed to recover for 3 weeks after surgery.

Ratiometric fiber photometry in the mPFC was conducted using an RZ5P processor running Synapse software (Tucker-Davis Technologies, Alachua, Florida, USA). A 405 nm LED (Doric Lenses, Quebec, Canada) and 470 nm LED (Doric Lenses) were modulated at 211 and 531 Hz to detect Ca^2+^-independent isosbestic signals and Ca^2+^-dependent signals, respectively. Light from the LEDs and GCaMP6s fluorescence was passed through a minicube (iFMC6, Doric Lenses) and the emitted light was detected using a fluorescence detector (DFD, Doric Lenses). Light power (10–30 μW) was measured at the tip of the fiber and adjusted using a light source device (LDFLS4, Doric Lenses). Fluorescence signals (1 kHz) were low-pass filtered with a frequency cutoff of 10 Hz and demodulated to 381 Hz using a MATLAB script. The time course of photobleaching was estimated by double exponential fitting to the fluorescence signals for the entire period, and photobleaching was corrected using a custom macro written in Igor Pro (WaveMetrics, Portland, OR, USA). The ΔF/F was calculated by dividing the change in the fluorescence signal by the baseline signal level.

### Immunohistochemistry and western blotting

Mice were deeply anesthetized with diethyl ether and transcardially perfused with heparinized (10 U/mL) phosphate-buffered saline (PBS), followed by PBS-buffered 4% (w/v) paraformaldehyde (PFA). Brains were removed, post-fixed in 4% PFA for 48 h at 4°C and cut into 100 μm coronal sections using a vibratome (VT1200S, Leica, Germany). The sections were post-fixed (1 h), permeabilized with 0.3% (v/v) Triton X-100 in PBS, and incubated in blocking buffer (5% normal goat serum, 5% horse serum, 5% donkey serum, and 0.5% BSA in PBS) for 2 h. Sections were successively incubated with primary [anti-mCherry (Abcam, Cat. # ab167453) and anti-GFP (Synaptic Systems, Cat. # 132 004); overnight at 4°C] and fluorescence (Cy3, Alexa fluor 647 or FITC: Jackson ImmunoResearch Laboratories, PA, USA) conjugated-secondary (3 h at room temperature) antibodies. Between each step, the sections were rinsed 3 times for 10 min with PBS. Images were acquired using an A1 confocal laser scanning microscope, and processed using the NIS Viewer (Nikon, Japan). Wide-field images of the entire brain section were acquired using an LSM 980 confocal microscope and Zeiss Application Suite ZEN (Zeiss, Germany).

For western blotting, mouse hippocampi or forebrains were homogenized in buffer (50 mM HEPES, 100 mM NaCl, 5 mM EGTA, 5 mM EDTA, and 1% Triton X-100, pH 7.4) containing a phosphatase inhibitor cocktail (GenDEPOT, TX, USA, Cat. # P3200) and proteinase inhibitor cocktail (Sigma-Aldrich, MO, USA, Cat. # P8340). To prepare subcellular hippocampal fractions, homogenates of the hippocampus were centrifuged at 1,200 × g for 10 min to remove nuclei and other large debris (P1). The supernatant (S1) was further centrifuged at 15,000 × g for 20 min to obtain a crude synaptosomal fraction (P2). All steps were performed using a synaptic protein extraction reagent (Syn-PER; Thermo Fisher Scientific) Proteins were separated by sodium dodecyl sulfate-polyacrylamide gel electrophoresis (SDS-PAGE) and transferred onto nitrocellulose membranes. The membranes were successively incubated with primary and horseradish peroxidase (HRP)-conjugated secondary (Jackson ImmunoResearch Laboratories, PA, USA) antibodies. Signals were detected using enhanced chemiluminescence (GE Healthcare, UK), and quantified using MetaMorph software (Molecular Devices, CA, USA). The following primary antibodies were purchased from commercial suppliers: anti-GluN1 (Cat. # 556308), anti-GluN2A (Cat. # 612286) and anti-GluN2B (Cat. # 610416) from BD Biosciences; anti-PSD95 (Cat. # MA1-045) from Invitrogen; anti-eEF2 (Cat. # 2332S), anti-p-eEF2 (Cat. # 2331S) and anti-GluN1 (Cat. # 5704S) from Cell Signaling Technology; α-tubulin (Cat. # T5168) from Sigma-Aldrich.

### Quantification and statistical analysis

Statistical analyses were performed using Igor Pro (WaveMetrics) and SPSS (IBM, Armonk, NY, USA). The normality of the collected data was determined using the Shapiro-Wilk test. The Mann-Whitney U test, Wilcoxon signed-rank test, or Kruskal-Wallis test was used to compare non-normally distributed samples. Samples that satisfied a normal distribution were compared using two-tailed Student’s *t*-tests. For multiple groups, a one-way ANOVA followed by Tukey’s HSD (honestly significant difference) post-hoc test was used to compare the samples. All bar graphs in the figures show the mean ± standard error of the mean (SEM). The levels of significance are indicated as follows: *P < 0.05, **P < 0.01, ***P < 0.001, and n.s., not significant (p ≥ 0.05). The number of cells, slices, or animals used for each experiment and statistical analyses are provided in the Supplementary Table 1.

## Acknowledgements

We thank Dr. Martin Horák and Anna Misiachna for helpful advice on NMDA current recordings in HEK293 cells; Young Sook Kim for genotyping of animals; Young Seon Cho for assistance in behavioral testing; Sung Pyo Oh for thoughtful discussion. We also thank Yoon Ju Kim for creating a visual summary of research findings. This study was supported by the National Research Foundation of Korea (NRF) grant funded by the Korea government (NRF-2017R1D1A1B03032935, 2018R1A5A2025964, 2017M3C7A1029609, and 2020R1A2C1014372).

## Author contributions

W.S.S, Y.-S.B and M.-H.K conceived the project and designed the experiments. W.S.S, Y.-S.B, and S.H.Y performed experiments and analyzed data. W.S.S and M.-H.K wrote the manuscript, with input from all other authors.

## Data availability

All data are available in the Source data or from the corresponding authors upon reasonable request.

