## Supplementary Fig for "Hyperactive GluN2B impairs neuroplasticity and cognition in phenylketonuria"

**Supplementary figures (Fig. 1−12)**

**Supplementary table 1**

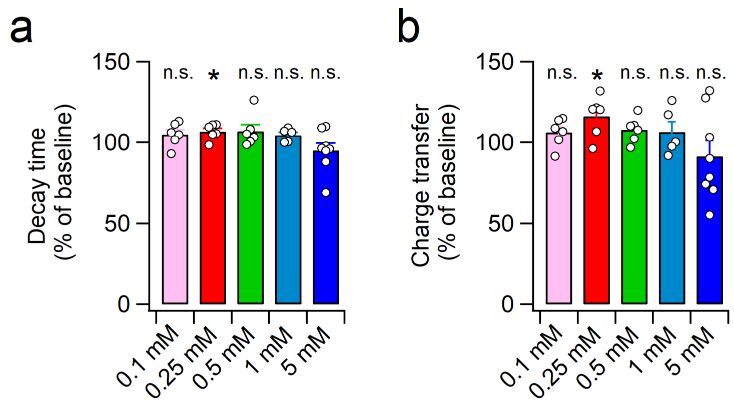


**Supplementary Fig. 1: The effects of different concentrations of L-Phe on the decay time and charge transfer of NMDA-EPSCs. a**, **b**, Decay time constant (**a**) and charge transfer (**b**) of NMDA-EPSCs during the L-Phe perfusion are summarized as a percentage of the baseline. NMDA-EPSCs were measured at −40 mV.

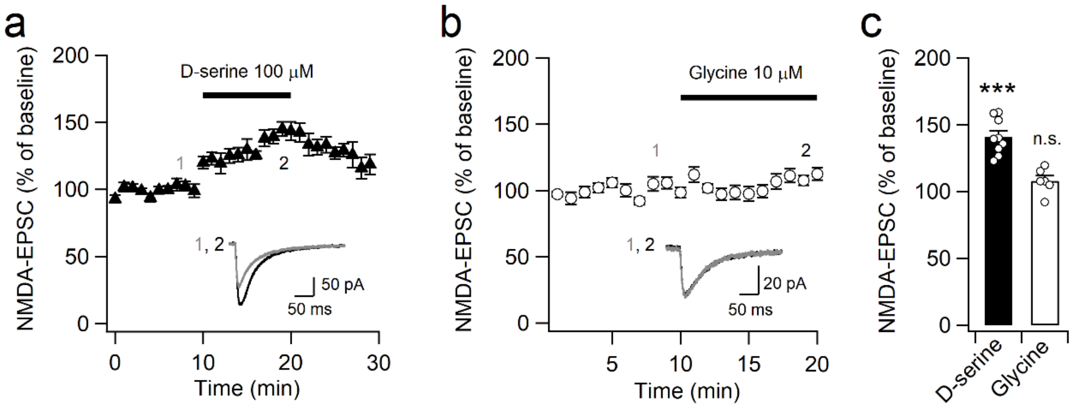


**Supplementary Fig. 2: NMDA-EPSCs at SC-CA1 synapse were increased by D-serine. a**, Application of 100 μM D-serine increases the peak amplitude of NMDA-EPSCs. Inset, sample trances of NMDA-EPSCs measured at the indicated time points (1, 2). **b**, Addition of glycine to ACSF did not affect NMDA-EPSCs. Time course of the peak amplitude of NMDA-EPSCs before and during bath application of 10 μM glycine. Inset, representative traces of NMDA-EPSCs obtained before and during perfusion of 10 μM glycine. **c**, Mean amplitudes of NMDA-EPSCs in the presence of D-serine or glycine in ACSF were compared with those obtained during the baseline recording.

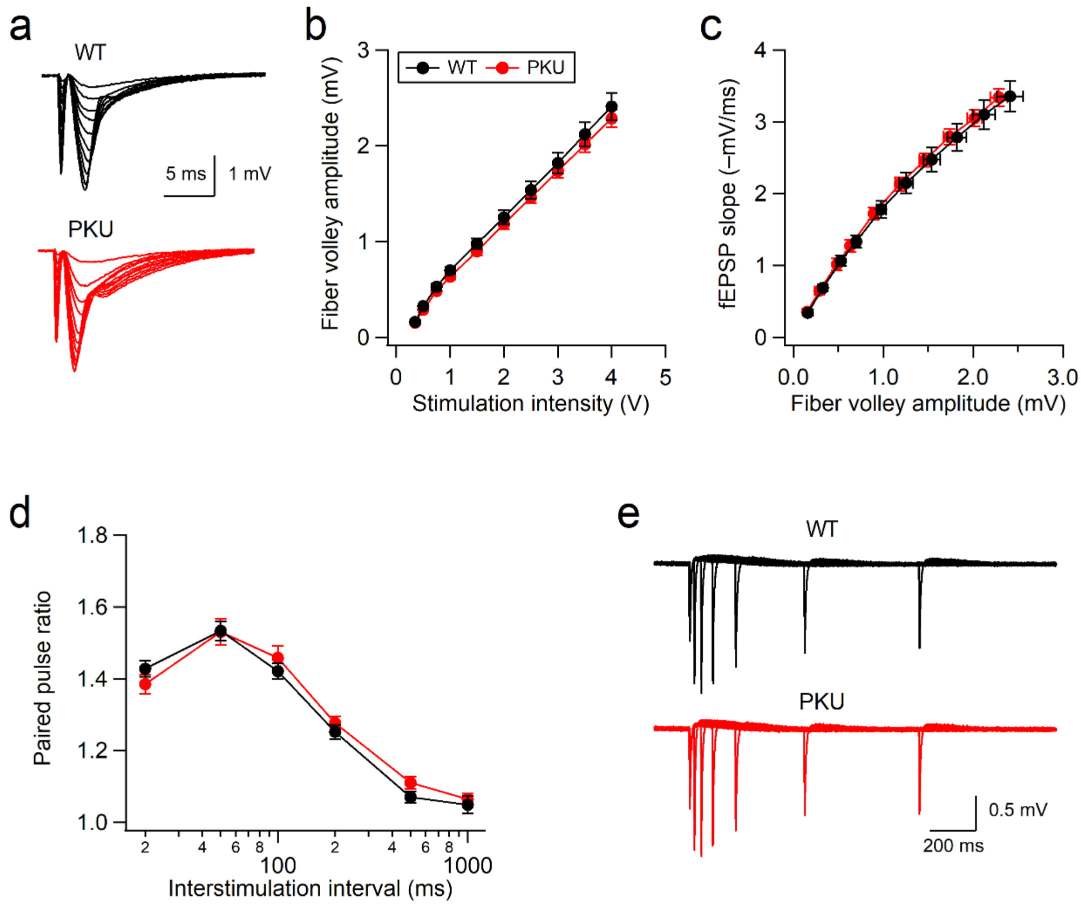


**Supplementary Fig. 3: Basal synaptic transmission at SC-CA1 synapse is normal in Pah$^{Enu2}$ mice**. **a**, fEPSPs were evoked in WT and Pah$^{Enu2}$ hippocampal slices with different stimulation intensities. **b**, The FV amplitude was plotted against stimulation intensity. **c**, The synaptic input-output ratio determined by plotting FV amplitude against fEPSP slope was not different between WT and Pah$^{Enu2}$ mice. **d**, Paired-pulse ratio of fEPSP slopes from WT and Pah$^{Enu2}$ slices are plotted as a function of the interstimulation interval. **e**, Sample traces of fEPSPs evoked by two consecutive stimuli with different interstimulus intervals.

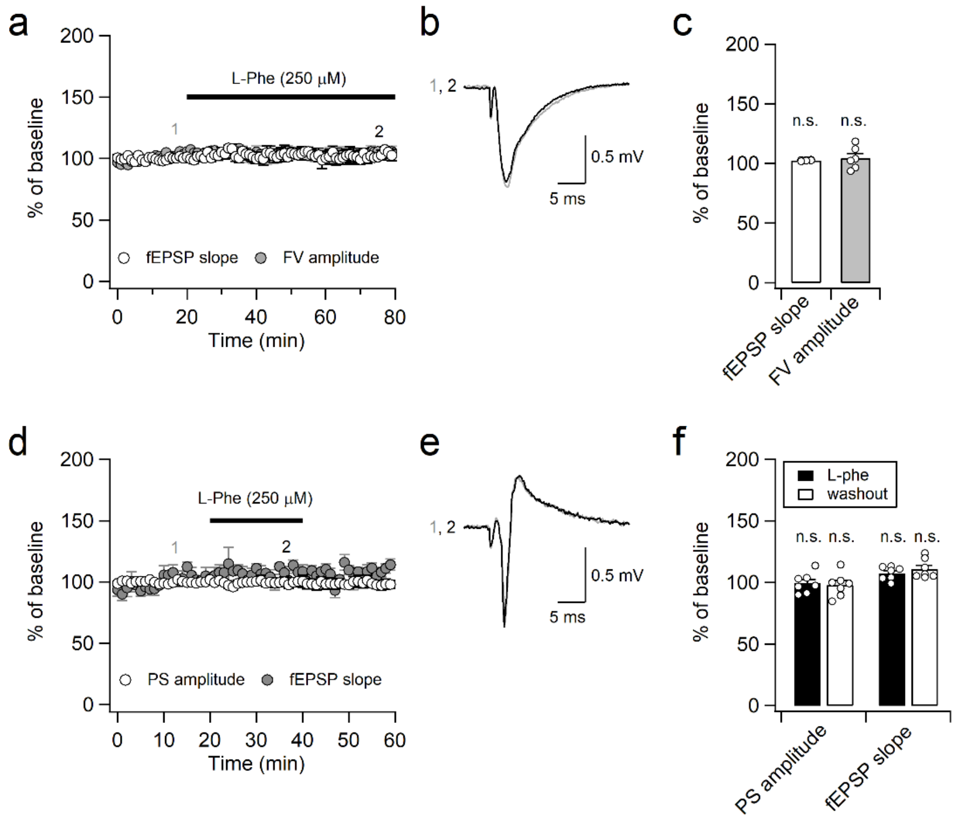


**Supplementary Fig. 4: L-Phe did not affect AMPAR-mediated synaptic transmission and population spikes (PS) in CA1 neurons. a**, The slopes of fEPSPs and the amplitude of fiber volley measured at the SC-CA1 synapse are plotted against the recording time. **b**, Sample fEPSP traces obtained before and during the L-Phe perfusion. **c**, The slopes of fEPSPs and the amplitudes of FV were not affected by L-Phe perfusion. **d**, The recording electrode was placed in the CA1 stratum pyramidale, and fEPSP and PS were induced by stimulating SC axons. The slopes of fEPSPs and the amplitudes of PSs were normalized to those obtained during baseline and plotted as a function of time. **e**, Sample fEPSP and PS traces obtained before and during the L-Phe perfusion. **f**, The slopes of fEPSPs and the amplitudes of PSs during and after L-Phe perfusion were summarized.

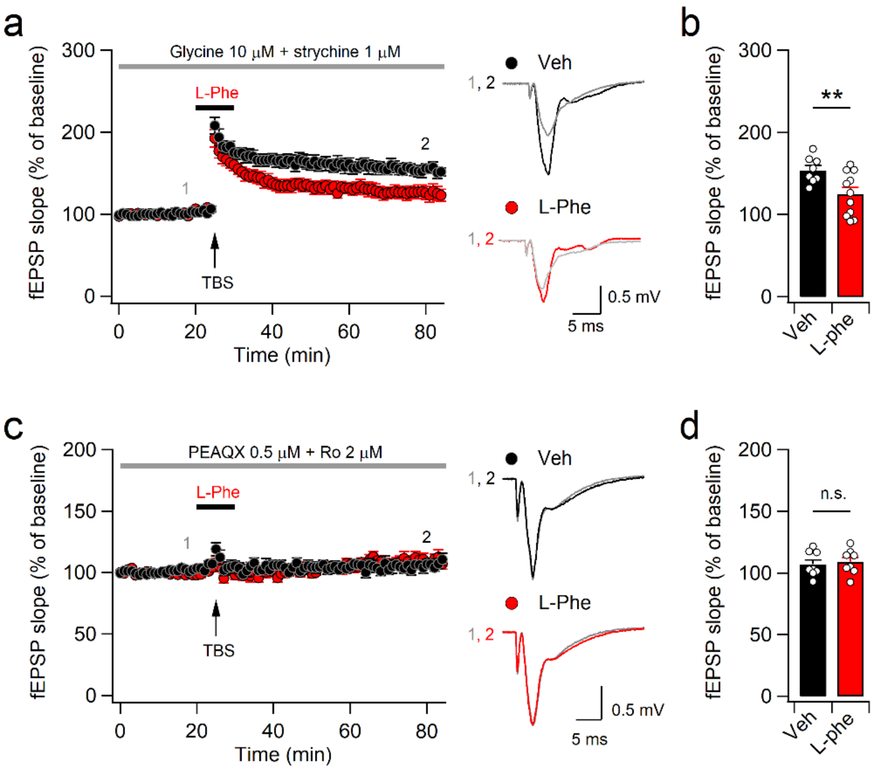


**Supplementary Fig. 5: L-Phe attenuates NMDAR-dependent LTP. a**, The slopes of fEPSPs measured in the presence of 10 µM glycine and the glycine receptor antagonist strychnine (1 µM) were plotted as a function of time. Right, sample traces obtained during baseline and the last 10 min of recording. **b**, The magnitude of LTP during the last 10 min of recording was significantly reduced by brief L-Phe perfusion during the peri-TBS period. **c**, TBS did not induce LTP in the presence of GluN2A and GluN2B inhibitors. Right, sample traces obtained before and after TBS. **d**, L-Phe and TBS had no effect on the slope of fEPSPs under GluN2A and GluN2B inhibition.

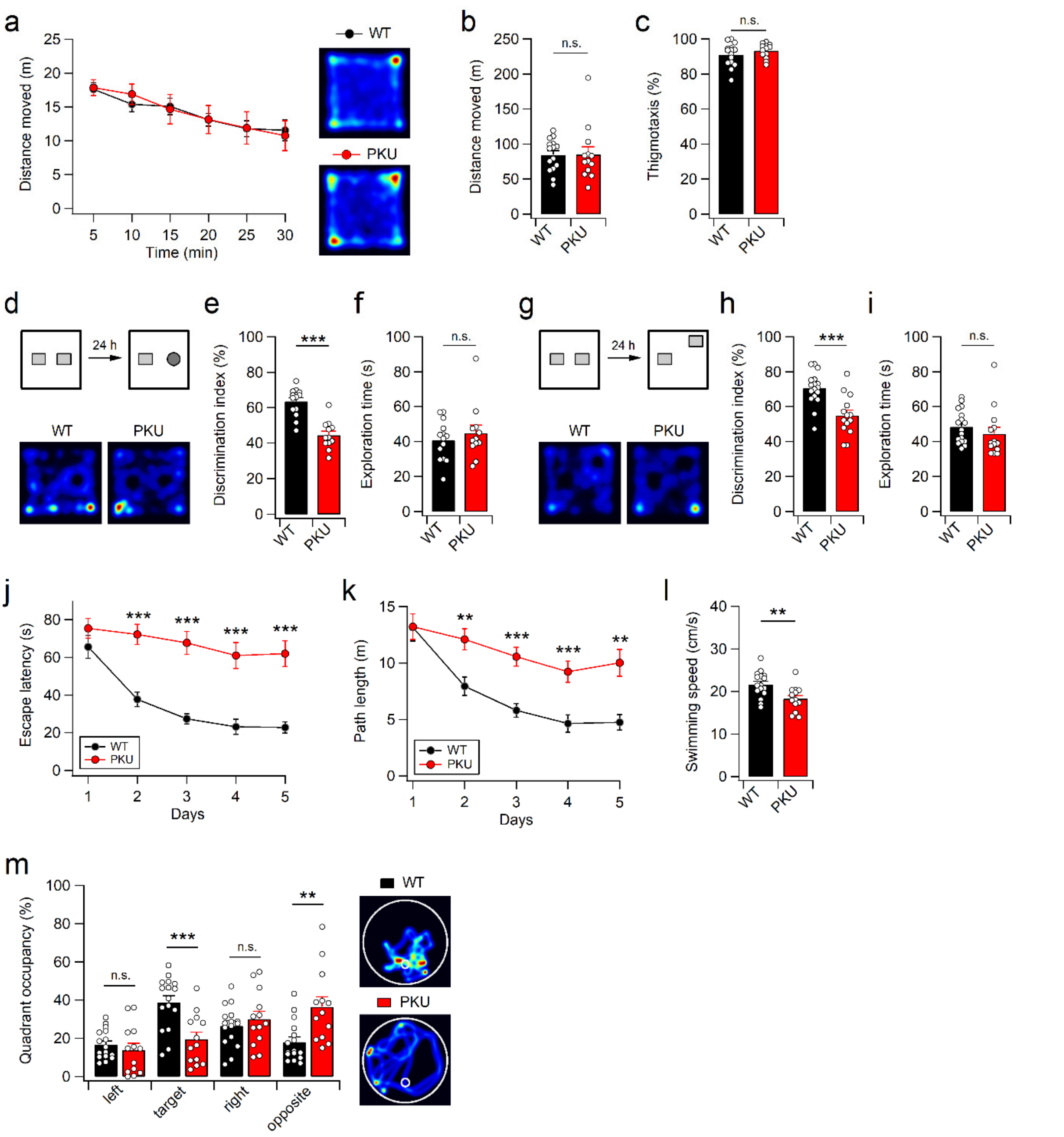


**Supplementary Fig. 6: Pah$^{Enu2}$ mice display impaired hippocampus-dependent learning and memory. a-c**, Normal open field activity of Pah$^{Enu2}$ mice. **a**, Activity of mice in the open-field box was plotted in 5 min bins for 30 min. Right, the representative activity path of mice during the first 10 min of the OFT test is shown. The distance moved of mice during the entire period (**b**) and time spent in the center during the first 15 min (**c**) are summarized. **d-i**, Pah$^{Enu2}$ mice display impaired NOR and OLM. **d**, Experimental design for NOR tests and representative activity path of mice during the NOR test session. Time spent exploring familiar and novel objects (**e**) and the discrimination index (**f**) for the NOR during the test session. **g-i**, Representative activity path (**g**), time spent exploring the two objects (**h**), and relative preference for the moved object (**i**) during the OLM test session. **j-m**, Impaired MWM performance of $Pah^{Enu2}$ mice. Escape latency (**j**) and swim distance (**k**) to find the hidden platform during the training period of MWM test. Swimming speed (**l**), quadrant occupancy (**m**, left) and representative swim path (**m**, right) of WT and $Pah^{Enu2}$ mice during the probe trials of MWM test.

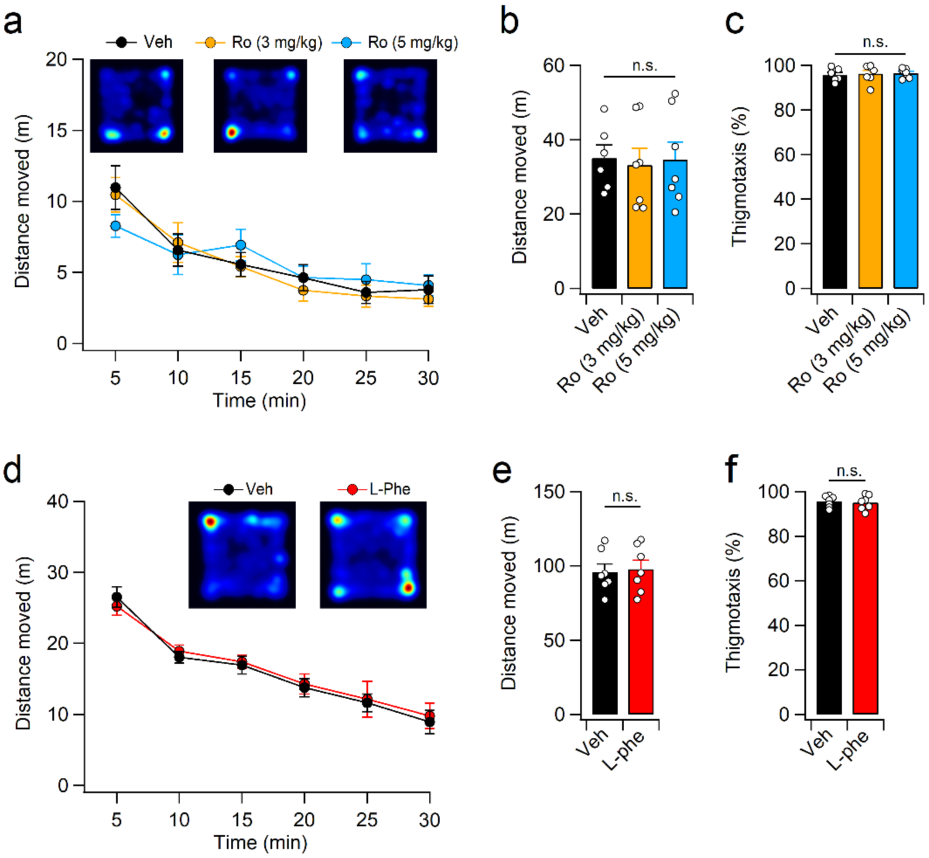


**Supplementary Fig. 7: Low concentrations of Ro and L-Phe did not affect open field activity of mice. a**, Activity of mice in the open-field box monitored for 30 min starting from 1 h after Ro administration (3 or 5 mg/kg, i.p.). **b**, Mice treated with 3 or 5 mg/kg Ro moved similar distances to those which received vehicle. **c**, Vehicle and Ro-treated mice spent similar amount of time in the outer region of the open field box. **d**, OFT was performed 30 min after L-Phe administration. (**e**, **f**) L-Phe challenge (1 mg/g, i.p.) did not affect distance moved (**e**) or thigmotactic behavior (**f**) of mice during the OFT.

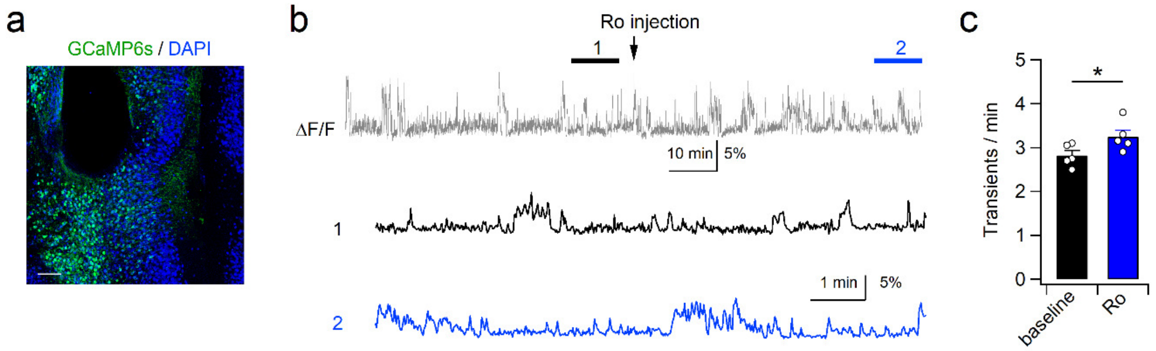


**Supplementary Fig. 8: Ro administration increases neural activity in the mPFC of Pah$^{Enu2}$ mice. a**, Immunofluorescence image showing GCaMP6s expression and cannula placement in the Pah$^{Enu2}$ mPFC. Calibration, 200 µm. **b**, Neural activity was monitored by fiber photometry. After 60 min of baseline recording, Pah$^{Enu2}$ mice were administered Ro (3 mg/kg, i.p.). **c**, The frequency of $Ca^{2+}$ transients before and after Ro administration is summarized.

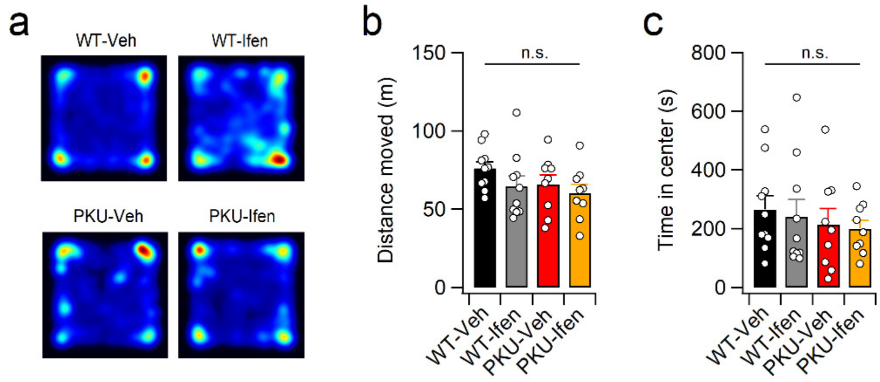


**Supplementary Fig. 9: Ifen did not affect the open field activity of WT and Pah$^{Enu2}$ mice.** **a**, Sample activity path of mice during the OFT. Mice were treated with ifen (5 mg/kg, i.p.) or vehicle 30 min before the OFT. **b**, **c**, Pretreatment of ifen had no effect on the distance moved (**b**) or time spent (**c**) in the center area during the OFT.

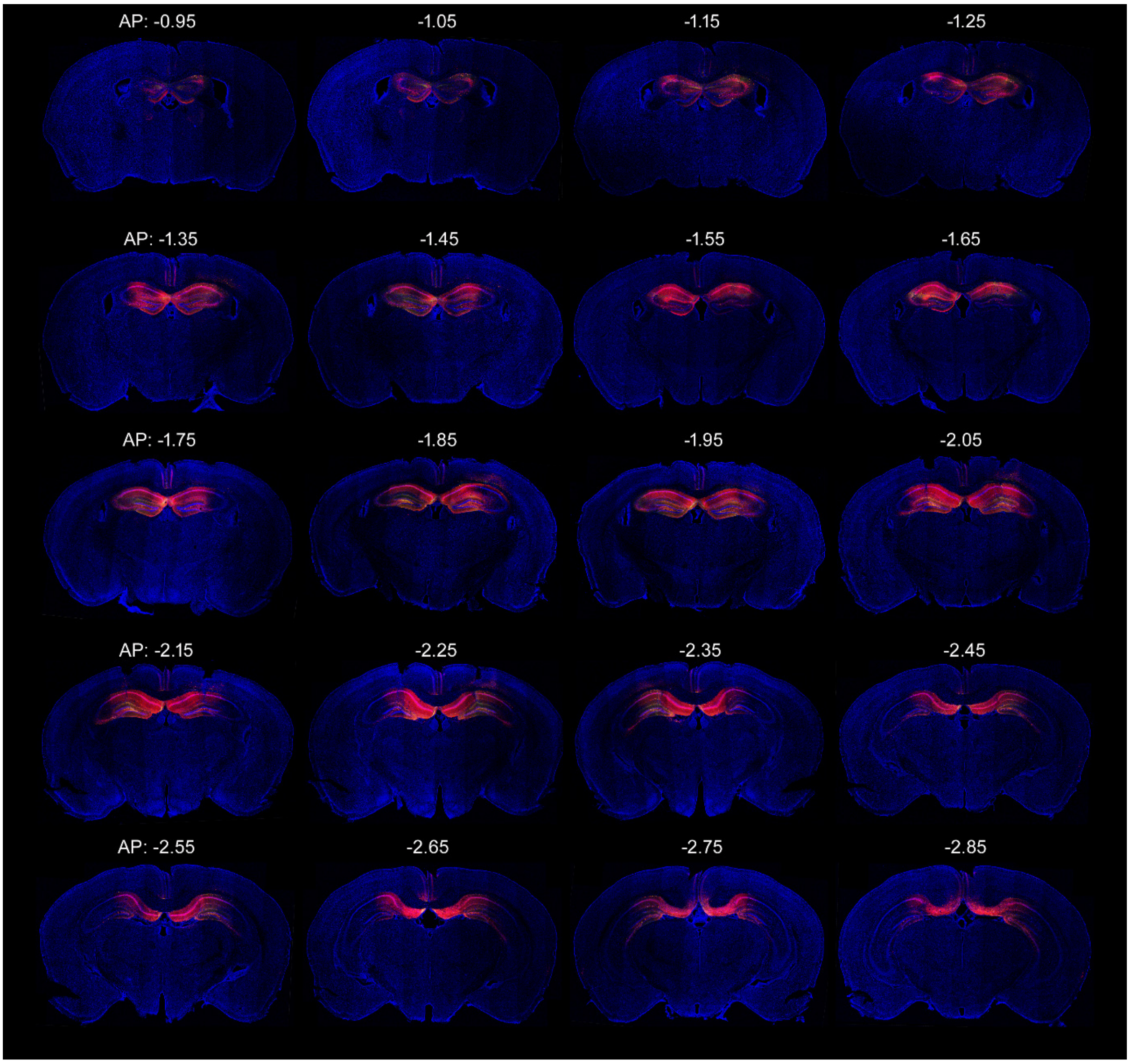


**Supplementary Fig. 10: A series of fluorescence images show the distribution of shGluN2B-expressing cells in the hippocampus of CaMKII-Cre;Pah$^{Enu2}$ mice.** Numbers represent coordinates of the anterior-posterior (AP) position from the bregma (mm). DAPI (blue) was used to identify brain regions and structures.

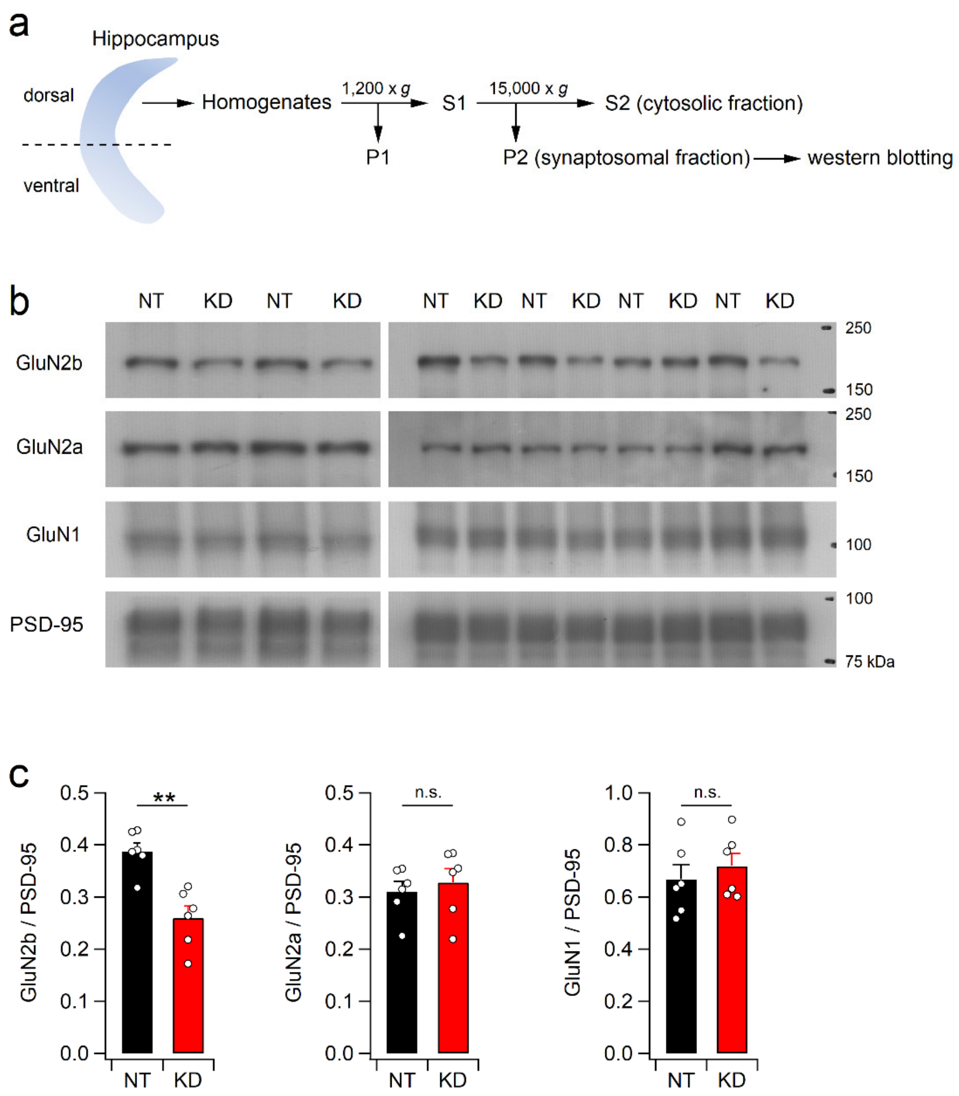


**Supplementary Fig. 11: Reduced GluN2B expression in the dorsal hippocampus of CaMKII-Cre mice expressing shGluN2B. a**, Diagram showing preparation of the synaptosomal fraction from the dorsal hippocampus. **b**, Expression levels of GluN1, GluN2A, and GluN2B were analyzed by western blotting. PSD-95 was used a loading control for normalization. NT and KD represent shNT and shGluN2B, respectively. **c**, CaMKII-Cre mice infected with AAV-shGluN2B exhibited reduced expression of GluN2B, not but GluN1 or GluN2A, in the synaptosomal fraction of dorsal hippocampus compared to those infected with AAV-shNT.

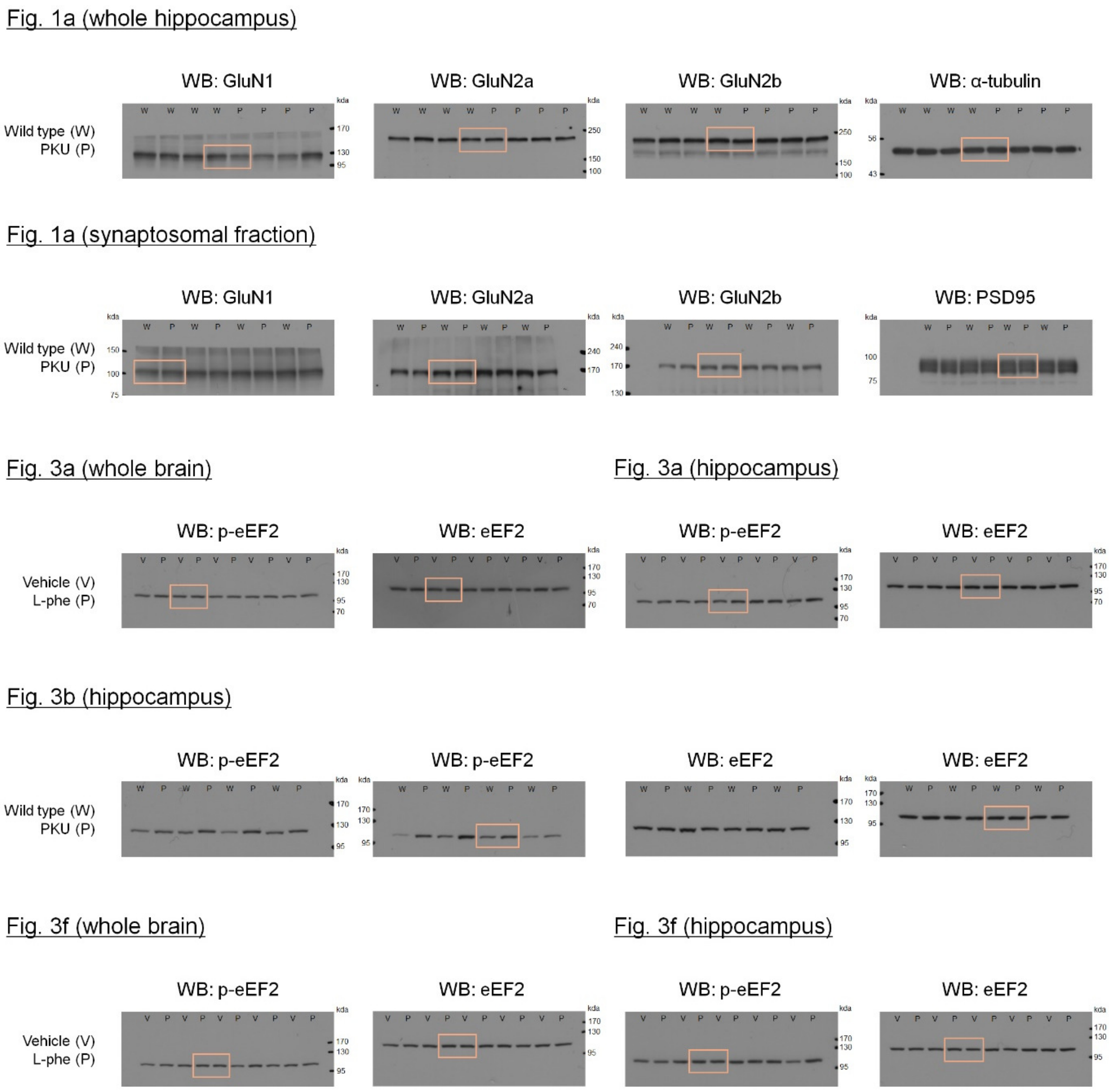


**Supplementary Fig. 12: Uncropped western blot images presented in the main figures.**

### Supplementary table 1. Statistical methods used for the analysis of experimental data in the study.

| | | | | Shapiro-Wilk test | | | Mann-Whitney U test<br>Wilkoxon signed-rank test | | | | Student's t-test | | | ANOVA<br>Kruskal-Wallis test | | Remark | |
|---|---|---|---|---|---|---|---|---|---|---|---|---|---|---|---|---|---|
| Figure | Experiment | Data number | Statistical method | W | p value | Normality | U | W | Z | p value | t | freedom | p value | F value<br>p value | post-hoc with Tukey's HSD<br>p value | | |
| Fig. 1a | evoked NMDA current amplitude | 0.1 mM: 6 cells from 2 mice | Paired t-test | diff(0.01 mM): 0.914 | diff(0.01 mM): p>0.05 | O | | | | | -0.903 | 5 | 0.408 | | | base vs. L-phe (0.01 mM) | |
| | | 0.25 mM: 6 cells from 4 mice | | diff(0.025 mM): 0.851 | diff(0.025 mM): p>0.05 | O | | | | | -1.048 | 5 | 0.343 | | | base vs. L-phe (0.025 mM) | |
| | | 0.5 mM: 6 cells from 2 mice | | diff(0.5 mM): 0.946 | diff(0.5 mM): p>0.05 | O | | | | | -1.608 | 5 | 0.169 | | | base vs. L-phe (0.5 mM) | |
| | | 1 mM: 5 cells from 2 mice | | diff(1 mM): 0.951 | diff(1 mM): p>0.05 | O | | | | | -0.453 | 4 | 0.674 | | | base vs. L-phe (1 mM) | |
| | | 5 mM: 10 cells from 3 mice | | diff(5 mM): 0.973 | diff(5 mM): p>0.05 | O | | | | | 1.182 | 9 | 0.268 | | | base vs. L-phe (5 mM) | |
| | | 0.1 mM: 6 cells from 2 mice<br>0.25 mM: 6 cells from 4 mice<br>0.5 mM: 6 cells from 2 mice<br>1 mM: 5 cells from 2 mice<br>5 mM: 10 cells from 3 mice | ANOVA | 0.1 mM: 0.910<br>0.25 mM: 0.814<br>0.5 mM: 0.912<br>1 mM: 0.949<br>5 mM: 0.962 | 0.1 mM: p>0.05<br>0.25 mM: p>0.05<br>0.5 mM: p>0.05<br>1 mM: p>0.05<br>5 mM: p>0.05 | O | | | | | | | | F(4,28)=1.683<br>p=0.182 | | | not shown in the figure |
| Fig. 1c | NMDA induced current ($I_{NMDA}$) | 0.1 mM: 6 cells from 2 mice | Paired t-test | diff(0.01 mM): 0.914 | diff(0.01 mM): p>0.05 | O | | | | | -2.916 | 5 | 0.0332 | | | NMDA vs. L-phe (0.01 mM) | |
| | | 0.25 mM: 11 cells from 6 mice | | diff(0.025 mM): 0.906 | diff(0.025 mM): p>0.05 | O | | | | | -4.365 | 10 | 0.00141 | | | NMDA vs. L-phe (0.025 mM) | |
| | | 0.5 mM: 6 cells from 3 mice | | diff(0.5 mM): 0.877 | diff(0.5 mM): p>0.05 | O | | | | | -4.815 | 5 | 0.00482 | | | NMDA vs. L-phe (0.5 mM) | |
| | | 1 mM: 8 cells from 4 mice | | diff(1 mM): 0.894 | diff(1 mM): p>0.05 | O | | | | | -5.036 | 7 | 0.00150 | | | NMDA vs. L-phe (1 mM) | |
| | | 5 mM: 5 cells from 3 mice | | diff(5 mM): 0.912 | diff(5 mM): p>0.05 | O | | | | | 4.659 | 4 | 0.00960 | | | NMDA vs. L-phe (5 mM) | |
| | | 10 mM: 3 cells from 2 mice | | dff(10 mM): 0.961 | dff(10 mM): p>0.05 | O | | | | | 5.162 | 2 | 0.0355 | | | NMDA vs. L-phe (10 mM) | |
| | | 0.1 mM: 6 cells from 2 mice<br>0.25 mM: 11 cells from 6 mice<br>0.5 mM: 6 cells from 3 mice<br>1 mM: 8 cells from 4 mice<br>5 mM: 5 cells from 3 mice<br>10 mM: 3 cells from 2 mice | ANOVA | 0.1 mM: 0.981<br>0.25 mM: 0.885<br>0.5 mM: 0.936<br>1 mM: 0.957<br>5 mM: 0.993<br>10 mM: 0.965 | 0.1 mM: p>0.05<br>0.25 mM: p>0.05<br>0.5 mM: p>0.05<br>1 mM: p>0.05<br>5 mM: p>0.05 | O | | | | | | | | F(5,33)=34.676<br>p=0.0000000000032142 | 0.277 | 0.25 mM vs. 0.1 mM | not shown in the figure |
| | | | | | | | | | | | | | | | 0.206 | 0.25 mM vs. 0.5 mM | |
| | | | | | | | | | | | | | | | 0.061 | 0.25 mM vs. 1 mM | |
| | | | | | | | | | | | | | | | 0.00000000202 | 0.25 mM vs. 5 mM | |
| | | | | | | | | | | | | | | | 0.0000000000361 | 0.25 mM vs. 10 mM | |
| Fig. 1e | NMDA induced current ($I_{NMDA}$) | PEAQX: 7 cells from 2 mice<br>Ro 25-6981: 9 cells from 2 mice<br>Ifenprodil: 8 cells from 2 mice | ANOVA | PEAQX: 0.922<br>Ro 25-6981: 0.960<br>Ifenprodil: 0.890 | PEAQX: p>0.05<br>Ro 25-6981: p>0.05<br>Ifenprodil: p>0.05 | O | | | | | | | | F(2,21)=16.769<br>p=0.000044 | 0.0000610 | PEAQX vs. Ro 25-6981 | |
| | | | | | | | | | | | | | | | 0.000398 | PEAQX vs. Ifenprodil | |
| | | | | | | | | | | | | | | | 0.766 | Ro 25-6981 vs. Ifenprodil | |
| Fig. 1f | evoked NMDA current amplitude | Veh: 9 cells from 5 mice<br>PEAQX: 9 cells from 5 mice<br>Ro 25-6981: 7 cells from 3 mice<br>Ifenprodil: 9 cells from 6 mice | ANOVA | Veh: 0.853<br>PEAQX: 0.869<br>Ro 25-6981: 0.949<br>Ifenprodil: 0.946 | Veh: p>0.05<br>PEAQX: p>0.05<br>Ro 25-6981: p>0.05<br>Ifenprodil: p>0.05 | O | | | | | | | | F(3,30)=10.109<br>p=0.000092 | 0.876 | Veh vs. PEAQX | |
| | | | | | | | | | | | | | | | 0.00347 | PEAQX vs. Ro 25-6981 | |
| | | | | | | | | | | | | | | | 0.0182 | PEAQX vs. Ifenprodil | |
| | | | | | | | | | | | | | | | 0.828 | Ro 25-6981 vs. Ifenprodil | |
| Fig. 1h | NMDA induced current ($I_{NMDA}$) | 8 cells | Paired t-test | diff: 0.972 | diff: p>0.05 | O | | | | | -3.453 | 7 | 0.0107 | | | base vs. L-phe | |
| Fig. 2b | Normalized expression level | Whole hippocampus<br>WT: 4 mice<br>PKU: 4 mice | Simple t-test | WT-GluN1: 0.981<br>PKU-GluN1: 0.992 | WT-GluN1: p>0.05<br>PKU-GluN1: p>0.05 | O | | | | | 4.685 | 6 | 0.00338 | | | WT vs. PKU | |
| | | | | WT-GluN2a: 0.896<br>PKU-GluN2a: 0.971 | WT-GluN2a: p>0.05<br>PKU-GluN2a: p>0.05 | O | | | | | 0.175 | 6 | 0.866 | | | WT vs. PKU | |
| | | | | WT-GluN2b: 0.950<br>PKU-GluN2b: 0.869 | WT-GluN1: p>0.05<br>PKU-GluN1: p>0.05 | O | | | | | -1.865 | 6 | 0.111 | | | WT vs. PKU | |
| | | Synaptosomal fraction<br>WT: 4 mice<br>PKU: 4 mice | | WT-GluN1: 0.812<br>PKU-GluN1: 0.818 | WT-GluN1: p>0.05<br>PKU-GluN1: p>0.05 | O | | | | | -1.839 | 6 | 0.116 | | | WT vs. PKU | |
| | | | | WT-GluN2a: 0.836<br>PKU-GluN2a: 0.960 | WT-GluN2a: p>0.05<br>PKU-GluN2a: p>0.05 | O | | | | | 0.323 | 6 | 0.758 | | | WT vs. PKU | |
| | | | | WT-GluN2b: 0.977<br>PKU-GluN2b: 0.886 | WT-GluN1: p>0.05<br>PKU-GluN1: p>0.05 | O | | | | | -0.53 | 6 | 0.615 | | | WT vs. PKU | |
| Fig. 2e | AMPA/NMDA ratio | WT: 11 cells from 4 mice<br>PKU: 9 cells from 3 mice | Simple t-test | WT: 0.925<br>PKU: 0.898 | WT: p>0.05<br>PKU: p>0.05 | O | | | | | -0.21 | 18 | 0.836 | | | WT vs. PKU | |
| Fig. 2g | NMDA induced current ($I_{NMDA}$) | WT: 8 cells from 4 mice<br>PKU: 9 cells from 4 mice | Simple t-test | WT-NMDA: 0.943<br>PKU-NMDA: 0.894 | WT-NMDA: p>0.05<br>PKU-NMDA: p>0.05 | O | | | | | -0.035 | 15 | 0.973 | | | WT vs. PKU | |
| | | | | WT-PEAQX: 0.940<br>PKU-PEAQX: 0.901 | WT-PEAQX: p>0.05<br>PKU-PEAQX: p>0.05 | O | | | | | 0.062 | 15 | 0.951 | | | WT vs. PKU | |
| | | | | WT-AP5: 0.939<br>PKU-AP5: 0.909 | WT-AP5: p>0.05<br>PKU-AP5: p>0.05 | O | | | | | -0.425 | 15 | 0.677 | | | WT vs. PKU | |
| Fig. 2h | TBS-induced LTP (fEPSP slope) | WT: 15 slices from 6 mice<br>PKU: 14 slices from 6 mice<br>WT-Phe: 13 slices from 4 mice<br>PKU-Phe: 13 slices from 3 mice | ANOVA | WT: 0.940<br>PKU: 0.960<br>WT-Phe: 0.981<br>PKU-Phe: 0.906 | WT: p>0.05<br>PKU: p>0.05<br>WT-Phe: p>0.05<br>PKU-Phe: p>0.05 | O | | | | | | | | F(3,51)=7.750<br>p=0.000233 | 0.988 | WT vs. PKU | |
| | | | | | | | | | | | | | | | 0.0124 | WT vs. WT-Phe | |
| | | | | | | | | | | | | | | | 0.00377 | PKU vs. PKU-Phe | |
| | | | | | | | | | | | | | | | 0.999 | WT-Phe vs. PKU-Phe | |
| Fig. 2i | TBS-induced LTP (fEPSP slope) | Veh-Veh: 12 slices from 10 mice<br>Veh-Phe: 11 slices from 9 mice<br>Ro-Phe: 10 slices from 7 mice<br>Ifen-Phe: 10 slices from 5 mice | Kruskal-Wallis test | Veh-Veh: 0.720<br>Veh-Phe: 0.806<br>Ro-Phe: 0.922<br>Ifen-Phe: 0.850 | Veh-Veh: p<0.05<br>Veh-Phe: p<0.05<br>Ro-Phe: p>0.05<br>Ifen-Phe: p>0.05 | X | | | | | | | | F(3,43)=15.126<br>p=0.001712 | 0.00115 | Veh-Veh vs. Veh-Phe | |
| | | | | | | | | | | | | | | | 0.00610 | Veh-Phe vs. Ro-Phe | |
| | | | | | | | | | | | | | | | 0.000774 | Veh-Phe vs. Ifen-Phe | |
| | | | | | | | | | | | | | | | 0.545 | Ro-Phe vs. Ifen-Phe | |
| Fig. 2k | TBS-induced LTP (fEPSP slope) | Veh: 12 slices from 4 mice<br>L-Phe: 10 slices from 4 mice | Simple t-test | Veh: 0.947<br>L-Phe: 0.959 | Veh: p>0.05<br>L-Phe: p>0.05 | O | | | | | 2.828 | 20 | 0.0104 | | | Veh vs. L-phe | |
| Fig. 2m | SP-LFS-induced LTD (fEPSP slope) | WT: 8 slices from 4 mice<br>PKU: 7 slices from 3 mice | Simple t-test | WT: 0.882<br>PKU: 0.941 | WT: p>0.05<br>PKU: p>0.05 | O | | | | | -0.783 | 13 | 0.448 | | | WT vs. PKU | |
| Fig. 2o | SP-LFS-induced LTD (fEPSP slope) | Veh-Veh: 9 slices from 6 mice<br>Veh-Phe: 9 slices from 7 mice<br>Ro-Phe: 9 slices from 4 mice | ANOVA | Veh-Veh: 0.934<br>Veh-Phe: 0.937<br>Ro-Phe: 0.991 | Veh-Veh: p>0.05<br>Veh-Phe: p>0.05<br>Ro-Phe: p>0.05 | O | | | | | | | | F(2,24)=5.029<br>p=0.014992 | 0.0350 | Veh-Veh vs. Veh-Phe | |
| | | | | | | | | | | | | | | | 0.987 | Veh-Veh vs. Ro-Phe | |
| | | | | | | | | | | | | | | | 0.0247 | Veh-Phe vs. Ro-Phe | |
| Fig. 3a | Normalized expression level | Veh(WB): 5 mice<br>Phe(WB): 5 mice | Simple t-test | Veh(WB): 0.958<br>Phe(WB): 0.955 | Veh(WB): p>0.05<br>Phe(WB): p>0.05 | O | | | | | -6.319 | 8 | 0.000228 | | | Veh vs. Phe | |
| | | Veh(HP): 5 mice<br>Phe(HP): 5 mice | | Veh(HP): 0.856<br>Phe(HP): 0.897 | Veh(HP): p>0.05<br>Phe(HP): p>0.05 | O | | | | | -5.438 | 8 | 0.00346 | | | Veh vs. Phe | |
| Fig. 3b | Normalized expression level | WT: 8 mice<br>PKU: 8 mice | Simple t-test | WT: 0.898<br>PKU: 0.957 | WT: p>0.05<br>PKU: p>0.05 | O | | | | | -5.155 | 14 | 0.000146 | | | WT vs. PKU | |
| Fig. 3e | Number of transients | 5 mice | Repeated measures ANOVA | diff[Baseline - L-Phe (1 h)]: 0.923<br>diff[L-Phe(1hr) - L-Phe (2 h)]: 0.984 | diff[Baseline - L-Phe (1 h)]: p>0.05<br>diff[L-Phe(1hr) - L-Phe (2 h)]: p>0.05 | O | | | | | | | | F(2,8)=40.551<br>p=0.000065 | 0.00702 | Baseline vs. L-Phe (1 h) | |
| | | | | | | | | | | | | | | | 0.00721 | Baseline vs. L-Phe (2 h) | |
| | | | | | | | | | | | | | | | 0.666 | L-Phe (1 h) vs. L-Phe (2 h) | |
| Fig. 3f | Normalized expression level | Ro-Veh(WB): 5 mice<br>Ro-Phe(WB): 5 mice | Simple t-test | Ro-Veh(WB): 0.828<br>Ro-Phe(WB): 0.886 | Ro-Veh(WB): p>0.05<br>Ro-Phe(WB): p>0.05 | O | | | | | 0.544 | 8 | 0.601 | | | Ro-Veh vs. Ro-Phe | |
| | | Ro-Veh(HP): 5 mice<br>Ro-Phe(HP): 5 mice | | Ro-Veh(HP): 0.868<br>Ro-Phe(HP): 0.950 | Ro-Veh(HP): p>0.05<br>Ro-Phe(HP): p>0.05 | O | | | | | -0.283 | 8 | 0.784 | | | Ro-Veh vs. Ro-Phe | |
| Fig. 3h | Number of transients | 5 mice | Repeated measures ANOVA | diff(baseline - L-phe): 0.899<br>diff (L-phe - Ro): 0.916 | diff(baseline - L-phe): p>0.05<br>diff (L-phe - Ro): p>0.05 | O | | | | | | | | F(2,8)=17.127<br>p=0.001285 | 0.0252 | baseline vs. L-phe | |
| | | | | | | | | | | | | | | | 0.245 | baseline vs. Ro | |
| | | | | | | | | | | | | | | | 0.0312 | L-phe vs. Ro | |
| Fig. 3j | Y-maze test (Spontaneous alternations) | Veh-Veh: 8 mice<br>Veh-Phe: 8 mice<br>Ro-Veh: 7 mice<br>Ro-Phe: 8 mice | ANOVA | Veh-Veh(%alteration): 0.944<br>Veh-Phe(%alteration): 0.835<br>Ro-Veh(%alteration): 0.821<br>Ro-Phe(%alteration): 0.908 | Veh-Veh(%alteration): p>0.05<br>Veh-Phe(%alteration): p>0.05<br>Ro-Veh(%alteration): p>0.05<br>Ro-Phe(%alteration): p>0.05 | O | | | | | | | | F(3,27)=0.995<br>p=0.410 | | | |
| Fig. 3l | Y-maze test (Novel arm exploration) | | | Veh-Veh(time): 0.851<br>Veh-Phe(time): 0.881<br>Ro-Veh(time): 0.920<br>Ro-Phe(time): 0.963 | Veh-Veh(time): p>0.05<br>Veh-Phe(time): p>0.05<br>Ro-Veh(time): p>0.05<br>Ro-Phe(time): p>0.05 | O | | | | | | | | F(3,27)=4.435<br>p=0.0117 | 0.0278 | Veh-Veh vs. Veh-Phe | |
| | | | | | | | | | | | | | | | 0.0485 | Veh-Phe vs. Ro-Veh | |
| | | | | | | | | | | | | | | | 0.0219 | Veh-Phe vs. Ro-Phe | |
| | | | | | | | | | | | | | | | 0.995 | Ro-Veh vs. Ro-Phe | |
| Fig. 3n | Novel object recognition | Veh-Veh: 8 mice<br>Veh-Phe: 8 mice<br>Ro-Veh: 8 mice<br>Ro-Phe: 8 mice | ANOVA | Veh-Veh(distance): 0.884<br>Veh-Phe(distance): 0.920<br>Ro-Veh(distance): 0.948<br>Ro-Phe(distance): 0.941 | Veh-Veh(distance): p>0.05<br>Veh-Phe(distance): p>0.05<br>Ro-Veh(distance): p>0.05<br>Ro-Phe(distance): p>0.05 | O | | | | | | | | F(3,28)=0.344<br>p=0.794 | | | |
| Fig. 3o | | | | Veh-Veh(time): 0.917<br>Veh-Phe(time): 0.879<br>Ro-Veh(time): 0.992<br>Ro-Phe(time): 0.936 | Veh-Veh(time): p>0.05<br>Veh-Phe(time): p>0.05<br>Ro-Veh(time): p>0.05<br>Ro-Phe(time): p>0.05 | O | | | | | | | | F(3,28)=0.016<br>p=0.997 | | | |
| Fig. 3p | | | | Veh-Veh(DI): 0.955<br>Veh-Phe(DI): 0.897<br>Ro-Veh(DI): 0.939<br>Ro-Phe(DI): 0.857 | Veh-Veh(DI): p>0.05<br>Veh-Phe(DI): p>0.05<br>Ro-Veh(DI): p>0.05<br>Ro-Phe(DI): p>0.05 | O | | | | | | | | F(3,28)=6.725<br>p=0.001471 | 0.0146 | Veh-Veh vs. Veh-Phe | |
| | | | | | | | | | | | | | | | 0.00155 | Veh-Phe vs. Ro-Veh | |
| | | | | | | | | | | | | | | | 0.0122 | Veh-Phe vs. Ro-Phe | |
| | | | | | | | | | | | | | | | 0.851 | Ro-Veh vs. Ro-Phe | |
| Fig. 4b | Novel object recognition | WT-Veh: 10 mice<br>WT-Ifen: 9 mice<br>PKU-Veh: 9 mice<br>PKU-Ifen: 9 mice | ANOVA | WT-Veh(time): 0.966<br>WT-Ifen(time): 0.873<br>PKU-Veh(time): 0.987<br>PKU-Ifen(time): 0.944 | WT-Veh(time): p>0.05<br>WT-Ifen(time): p>0.05<br>PKU-Veh(time): p>0.05<br>PKU-Ifen(time): p>0.05 | O | | | | | | | | F(3,33)=0.657<br>p=0.584 | | | |
| Fig. 4c | | | | WT-Veh(DI): 0.944<br>WT-Ifen(DI): 0.974<br>PKU-Veh(DI): 0.972<br>PKU-Ifen(DI): 0.956 | WT-Veh(DI): p>0.05<br>WT-Ifen(DI): p>0.05<br>PKU-Veh(DI): p>0.05<br>PKU-Ifen(DI): p>0.05 | O | | | | | | | | F(3,33)=23.191<br>p=0.000000029234 | 0.728 | WT-Veh vs. WT-Ifen | |
| | | | | | | | | | | | | | | | 0.0000000742 | WT-Veh vs. PKU-Veh | |
| | | | | | | | | | | | | | | | 0.00000100 | PKU-Veh vs. PKU-Ifen | |
| | | | | | | | | | | | | | | | 0.829 | PKU-Ifen vs. WT-Veh | |
| Fig. 4e | Novel object recognition | WT-Veh: 9 mice<br>PKU-Veh: 6 mice<br>PKU-Ro: 6 mice | ANOVA | WT-Veh(time): 0.919<br>PKU-Veh(time): 0.963<br>PKU-Ro(time): 0.882 | WT-Veh(time): p>0.05<br>PKU-Veh(time): p>0.05<br>PKU-Ro(time): p>0.05 | O | | | | | | | | F(2,18)=0.647<br>p=0.535 | | | |
| Fig. 4f | | | | WT-Veh(DI): 0.962<br>PKU-Veh(DI): 0.942<br>PKU-Ro(DI): 0.957 | WT-Veh(DI): p>0.05<br>PKU-Veh(DI): p>0.05<br>PKU-Ro(DI): p>0.05 | O | | | | | | | | F(2,18)=14.612<br>p=0.000170 | 0.000125 | WT-Veh vs. PKU-Veh | |
| | | | | | | | | | | | | | | | 0.3580000000 | WT-Veh vs. PKU-Ro | |
| | | | | | | | | | | | | | | | 0.00557000 | PKU-Veh vs. PKU-Ro | |
| Fig. 4h | Object location memory | WT-Veh: 10 mice<br>WT-Ifen: 10 mice<br>PKU-Veh: 9 mice<br>PKU-Ifen: 9 mice | ANOVA | WT-Veh(time): 0.851<br>WT-Ifen(time): 0.881<br>PKU-Veh(time): 0.963<br>PKU-Ifen(time): 0.920 | WT-Veh(time): p>0.05<br>WT-Ifen(time): p>0.05<br>PKU-Veh(time): p>0.05<br>PKU-Ifen(time): p>0.05 | O | | | | | | | | F(3,34)=0.205<br>p=0.892 | | | |
| Fig. 4i | | | | WT-Veh(DI): 0.951<br>WT-Ifen(DI): 0.963<br>PKU-Veh(DI): 0.917<br>PKU-Ifen(DI): 0.959 | WT-Veh(DI): p>0.05<br>WT-Ifen(DI): p>0.05<br>PKU-Veh(DI): p>0.05<br>PKU-Ifen(DI): p>0.05 | O | | | | | | | | F(3,34)=17.616<br>p=0.00000044988 | 0.760 | WT-Veh vs. WT-Ifen | |
| | | | | | | | | | | | | | | | 0.00000100 | WT-Veh vs. PKU-Veh | |
| | | | | | | | | | | | | | | | 0.0000130 | PKU-Veh vs. PKU-Ifen | |
| | | | | | | | | | | | | | | | 0.899 | PKU-Ifen vs. WT-Veh | |
| Fig. 4m | NMDA induced current ($I_{NMDA}$) | uninfected: 6 cells from 4 mice<br>shNT: 9 cells from 2 mice<br>shGluN2B: 11 cells from 3 mice | ANOVA | uninfected: 0.988<br>shNT: 0.868<br>shGluN2B: 0.878 | uninfected: p>0.05<br>shNT: p>0.05<br>shGluN2B: p>0.05 | O | | | | | | | | F(2,23)=33.138<br>p=0.00000016842 | 0.165 | uninfected vs. shNT | |
| | | | | | | | | | | | | | | | 0.000000503 | uninfected vs. shGluN2B | |
| | | | | | | | | | | | | | | | 0.000009 | shNT vs. shGluN2B | |

| | | | | | | | | | | | | | | | | | |
|---|---|---|---|---|---|---|---|---|---|---|---|---|---|---|---|---|---|
| Fig. 4n | Escape latency (MWM - training session) | WT-NT: 9 mice<br>WT-KD: 10 mice<br>PKU-NT: 10 mice<br>PKU-KD: 11 mice | 2-way ANOVA | | | | | | | | | | | F(3,180)=15.212<br>p=0.0000000072<br>625 | 3.87E-08 | WT-NT vs. PKU-NT | |
| | | | | | | | | | | | | | | | 9.0102E-07 | WT-KD vs. PKU-NT | |
| | | | | | | | | | | | | | | | 0.000131 | PKU-NT vs. PKU-KD | |
| | | | | | | | | | | | | | | | 0.192 | PKU-KD vs. PKU-NT | |
| Fig. 4p | Swimming speed (MWM - probe test) | WT-Scr: 9 mice<br>WT-2bKD: 10 mice<br>PKU-Scr: 10 mice<br>PKU-2bKD: 11 mice | ANOVA | WT-Scr: 0.910<br>WT-2bKD: 0.837<br>PKU-Scr: 0.904<br>PKU-2bKD: 0.891 | WT-Scr: p>0.05<br>WT-2bKD: p>0.05<br>PKU-Scr: p>0.05<br>PKU-2bKD: p>0.05 | O | | | | | | | | F(3,36)=0.511<br>p=0.677 | | | |
| Fig. 4q | Quadrant occupancy (MWM - probe test) | WT-NT: 9 mice<br>WT-KD: 10 mice<br>PKU-NT: 10 mice<br>PKU-KD: 11 mice | ANOVA | WT-NT(Opp): 0.975<br>WT-NT(Right): 0.957<br>WT-NT(Target): 0.936<br>WT-NT(Left): 0.902 | WT-NT(Opp): p>0.05<br>WT-NT(Right): p>0.05<br>WT-NT(Target): p>0.05<br>WT-NT(Left): p>0.05 | O | | | | | | | | F(3,32)=16.37<br>p=0.000001 | 0.000012 | Target vs. Opp | |
| | | | | | | | | | | | | | | | 0.00045 | Target vs. Right | |
| | | | | | | | | | | | | | | | 0.000003 | Target vs. Left | |
| | | | | WT-KD(Opp): 0.927<br>WT-KD(Right): 0.937<br>WT-KD(Target): 0.956<br>WT-KD(Left): 0.984 | WT-KD(Opp): p>0.05<br>WT-KD(Right): p>0.05<br>WT-KD(Target): p>0.05<br>WT-KD(Left): p>0.05 | O | | | | | | | | F(3,36)=9.759<br>p=0.000075 | 0.00136 | Target vs. Opp | |
| | | | | | | | | | | | | | | | 0.000435 | Target vs. Right | |
| | | | | | | | | | | | | | | | 0.000246 | Target vs. Left | |
| | | | | PKU-NT(Opp): 0.934<br>PKU-NT(Right): 0.949<br>PKU-NT(Target): 0.977<br>PKU-NT(Left): 0.982 | PKU-NT(Opp): p>0.05<br>PKU-NT(Right): p>0.05<br>PKU-NT(Target): p>0.05<br>PKU-NT(Left): p>0.05 | O | | | | | | | | F(3,36)=2.442<br>p=0.0800 | 0.606 | Target vs. Opp | |
| | | | | | | | | | | | | | | | 0.996 | Target vs. Right | |
| | | | | | | | | | | | | | | | 0.484 | Target vs. Left | |
| | | | | PKU-KD(Opp): 0.965<br>PKU-KD(Right): 0.973<br>PKU-KD(Target): 0.939<br>PKU-KD(Left): 0.987 | PKU-KD(Opp): p>0.05<br>PKU-KD(Right): p>0.05<br>PKU-KD(Target): p>0.05<br>PKU-KD(Left): p>0.05 | O | | | | | | | | F(3,40)=12.759<br>p=0.000005 | 0.000023 | Target vs. Opp | |
| | | | | | | | | | | | | | | | 0.00444 | Target vs. Right | |
| | | | | | | | | | | | | | | | 0.00002 | Target vs. Left | |
| | | | | WT-NT(Target): 0.936<br>WT-KD(Target): 0.956<br>PKU-NT(Target): 0.977<br>PKU-KD(Target): 0.939 | WT-NT(Target): p>0.05<br>WT-KD(Target): p>0.05<br>PKU-NT(Target): p>0.05<br>PKU-KD(Target): p>0.05 | O | | | | | | | | F(3,36)=5.344<br>p=0.003783 | 0.00736 | PKU-NT vs. WT-NT | |
| | | | | | | | | | | | | | | | 0.00888 | PKU-NT vs. WT-KD | |
| | | | | | | | | | | | | | | | 0.0459 | PKU-NT vs. PKU-KD | |
| Supplementary Fig. 1a | evoked NMDA current (decay time) | 0.1 mM: 6 cells from 2 mice | Paired t-test | diff(0.01 mM): 0.937 | diff(0.01 mM): p>0.05 | O | | | | | -1.873 | 5 | 0.120 | | | base vs. L-phe (0.01 mM) | |
| | | 0.25 mM: 6 cells from 4 mice | | diff(0.025 mM): 0.955 | diff(0.025 mM): p>0.05 | O | | | | | -3.319 | 5 | 0.021 | | | base vs. L-phe (0.025 mM) | |
| | | 0.5 mM: 6 cells from 2 mice | Wilkoxon signed-rank test | diff(0.5 mM): 0.767 | diff(0.5 mM): p<0.05 | X | | | -1.782 | 0.075 | | | | | | base vs. L-phe (0.5 mM) | |
| | | 1 mM: 5 cells from 2 mice | Paired t-test | diff(1 mM): 0.881 | diff(1 mM): p>0.05 | O | | | | | -2.458 | 4 | 0.070 | | | base vs. L-phe (1 mM) | |
| | | 5 mM: 8 cells from 3 mice | | diff(5 mM): 0.855 | diff(5 mM): p>0.05 | O | | | | | 1.187 | 7 | 0.274 | | | base vs. L-phe (5 mM) | |
| | | 0.1 mM: 6 cells from 2 mice<br>0.25 mM: 6 cells from 4 mice<br>0.5 mM: 6 cells from 2 mice<br>1 mM: 5 cells from 2 mice<br>5 mM: 8 cells from 3 mice | ANOVA | 0.1 mM: 0.950<br>0.25 mM: 0.880<br>0.5 mM: 0.801<br>1 mM: 0.867<br>5 mM: 0.885 | 0.1 mM: p>0.05<br>0.25 mM: p>0.05<br>0.5 mM: p>0.05<br>1 mM: p>0.05<br>5 mM: p>0.05 | O | | | | | | | | F(4,26)=2.206<br>p=0.096 | | | not shown in the figure |
| Supplementary Fig. 1b | evoked NMDA current (charge stransfer) | 0.1 mM: 6 cells from 2 mice | Paired t-test | diff(0.01 mM): 0.865 | diff(0.01 mM): p>0.05 | O | | | | | -1.478 | 5 | 0.200 | | | base vs. L-phe (0.01 mM) | |
| | | 0.25 mM: 6 cells from 4 mice | | diff(0.025 mM): 0.963 | diff(0.025 mM): p>0.05 | O | | | | | -2.983 | 5 | 0.031 | | | base vs. L-phe (0.025 mM) | |
| | | 0.5 mM: 6 cells from 2 mice | | diff(0.5 mM): 0.942 | diff(0.5 mM): p>0.05 | O | | | | | -2.178 | 5 | 0.081 | | | base vs. L-phe (0.5 mM) | |
| | | 1 mM: 5 cells from 2 mice | | diff(1 mM): 0.911 | diff(1 mM): p>0.05 | O | | | | | -0.933 | 4 | 0.404 | | | base vs. L-phe (1 mM) | |
| | | 5 mM: 8 cells from 3 mice | | diff(5 mM): 0.967 | diff(5 mM): p>0.05 | O | | | | | 1.123 | 7 | 0.298 | | | base vs. L-phe (5 mM) | |
| | | 0.1 mM: 6 cells from 2 mice<br>0.25 mM: 6 cells from 4 mice<br>0.5 mM: 6 cells from 2 mice<br>1 mM: 5 cells from 2 mice<br>5 mM: 8 cells from 3 mice | ANOVA | 0.1 mM: 0.919<br>0.25 mM: 0.917<br>0.5 mM: 0.972<br>1 mM: 0.908<br>5 mM: 0.930 | 0.1 mM: p>0.05<br>0.25 mM: p>0.05<br>0.5 mM: p>0.05<br>1 mM: p>0.05<br>5 mM: p>0.05 | O | | | | | | | | F(4,26)=1.952<br>p=0.132 | | | not shown in the figure |
| Supplementary Fig. 2c | evoked NMDA current amplitude | 9 cells from 3 mice | Paired t-test | diff: 0.911 | diff: p>0.05 | O | | | | | -7.574 | 8 | 0.0000650 | | | base vs. D-serine | |
| | | 6 cells from 3 mice | Paired t-test | diff: 0.925 | diff: p>0.05 | O | | | | | -2.001 | 5 | 0.102 | | | base vs. Glycine | |
| Supplementary Fig. 3c | fEPSP slope | 6 slices from 3 mice | Simple t-test | Base: 0.958<br>L-Phe: 0.958 | Base: p>0.05<br>L-Phe: p>0.05 | O | | | | | -1.199 | 10 | 0.258 | | | Base vs. L-Phe | |
| | Fiber volley amplitude | 6 slices from 3 mice | Simple t-test | Base: 0.930<br>L-Phe: 0.954 | Base: p>0.05<br>L-Phe: p>0.05 | O | | | | | -0.492 | 10 | 0.643 | | | Base vs. L-Phe | |
| Supplementary Fig. 3f | fEPSP slope | 7 slices from 2 mice | ANOVA | Base: 0.944<br>L-Phe: 0.903<br>WO: 0.831 | Base: p>0.05<br>L-Phe: p>0.05<br>WO: p>0.05 | O | | | | | | | | F(2,18)=2.45<br>p=0.115 | | | |
| | Population spike | 7 slices from 2 mice | ANOVA | Base: 0.898<br>L-Phe: 0.939<br>WO: 0.958 | Base: p>0.05<br>L-Phe: p>0.05<br>WO: p>0.05 | O | | | | | | | | F(2,18)=0.13<br>p=0.879 | | | |
| Supplementary Fig. 4d | Paired pulse ratio (20 ms interval) | WT: 17 cells from 5 mice<br>PKU: 14 cells from 4 mice | Simple t-test | WT: 0.967<br>PKU: 0.947 | WT: p>0.05<br>PKU: p>0.05 | O | | | | | 1.222 | 29 | 0.232 | | | WT vs. PKU | not shown in the figure |
| | Paired pulse ratio (50 ms interval) | | Simple t-test | WT: 0.915<br>PKU: 0.964 | WT: p>0.05<br>PKU: p>0.05 | O | | | | | 0.071 | 29 | 0.944 | | | WT vs. PKU | |
| | Paired pulse ratio (100 ms interval) | | Simple t-test | WT: 0.983<br>PKU: 0.940 | WT: p>0.05<br>PKU: p>0.05 | O | | | | | -0.93 | 22.314 | 0.362 | | | WT vs. PKU | |
| | Paired pulse ratio (200 ms interval) | | Simple t-test | WT: 0.951<br>PKU: 0.976 | WT: p>0.05<br>PKU: p>0.05 | O | | | | | -0.905 | 29 | 0.373 | | | WT vs. PKU | |
| | Paired pulse ratio (500 ms interval) | | Simple t-test | WT: 0.966<br>PKU: 0.884 | WT: p>0.05<br>PKU: p>0.05 | O | | | | | -1.714 | 29 | 0.0971 | | | WT vs. PKU | |
| | Paired pulse ratio (1000 ms interval) | | Mann-Whitney U test | WT: 0.823<br>PKU: 0.915 | WT: p<0.05<br>PKU: p>0.05 | X | 90 | 243 | -1.151 | 0.262 | | | | | | WT vs. PKU | |
| Supplementary Fig. S5b | TBS-induced LTP (fEPSP slope) | Veh: 8 slices from 4 mice<br>L-Phe: 11 slices from 4 mice | Simple t-test | Veh: 0.966<br>L-Phe: 0.903 | Veh: p>0.05<br>L-Phe: p>0.05 | O | | | | | 2.995 | 17 | 0.00830 | | | Veh vs. L-Phe | |
| Supplementary Fig. 5d | TBS-induced LTP (fEPSP slope) | Veh: 8 slices from 3 mice<br>L-Phe: 8 slices from 3 mice | Simple t-test | Veh: 0.899<br>L-Phe: 0.968 | Veh: p>0.05<br>L-Phe: p>0.05 | O | | | | | -0.407 | 14 | 0.690 | | | Veh vs. L-Phe | |
| Supplementary Fig. 6b | OFT (total distance moved) | WT: 15 mice<br>PKU: 13 mice | Mann-Whitney U test | WT: 0.956<br>PKU: 0.822 | WT: p>0.05<br>PKU: p<0.05 | X | 85 | 176 | -0.576 | 0.586 | | | | | | WT vs. PKU | |
| Supplementary Fig. 6c | OFT (thigmotaxis) | | Simple t-test | WT: 0.949<br>PKU: 0.925 | WT: p>0.05<br>PKU: p>0.05 | O | | | | | -1.029 | 26 | 0.313 | | | WT vs. PKU | |
| Supplementary Fig. 6e | NOR (discrimination index) | WT: 14 mice<br>PKU: 12 mice | Simple t-test | WT: 0.919<br>PKU: 0.966 | WT: p>0.05<br>PKU: p>0.05 | O | | | | | 6.145 | 24 | 0.00000200 | | | WT vs. PKU | |
| Supplementary Fig. 6f | NOR (exploration time) | | Mann-Whitney U test | WT: 0.959<br>PKU: 0.822 | WT: p>0.05<br>PKU: p<0.05 | X | 78 | 183 | -0.309 | 0.781 | | | | | | WT vs. PKU | |
| Supplementary Fig. 6h | OLM (discrimination index) | WT: 17 mice<br>PKU: 14 mice | Simple t-test | WT: 0.943<br>PKU: 0.962 | WT: p>0.05<br>PKU: p>0.05 | O | | | | | 4.111 | 29 | 0.000296 | | | WT vs. PKU | |
| Supplementary Fig. 6i | OLM (exploration time) | | Mann-Whitney U test | WT: 0.926<br>PKU: 0.768 | WT: p>0.05<br>PKU: p<0.05 | X | 74 | 179 | -1.786 | 0.0771 | | | | | | WT vs. PKU | |
| Supplementary Fig. 6j | Escape latency (MWM - training day 1) | WT: 16 mice<br>PKU: 13 mice | Mann-Whitney U test | WT: 0.860<br>PKU: 0.741 | WT: p<0.05<br>PKU: p<0.05 | X | 81 | 217 | -1.037 | 0.329 | | | | | | WT vs. PKU | |
| | Escape latency (MWM - training day 2) | | Mann-Whitney U test | WT: 0.956<br>PKU: 0.849 | WT: p>0.05<br>PKU: p<0.05 | X | 14.5 | 150.5 | -3.935 | 0.0000150 | | | | | | WT vs. PKU | |
| | Escape latency (MWM - training day 3) | | Simple t-test | WT: 0.950<br>PKU: 0.901 | WT: p>0.05<br>PKU: p>0.05 | O | | | | | -6.024 | 17.02 | 0.0000140 | | | WT vs. PKU | |
| | Escape latency (MWM - training day 4) | | Mann-Whitney U test | WT: 0.854<br>PKU: 0.927 | WT: p<0.05<br>PKU: p>0.05 | X | 22 | 158 | -3.599 | 0.000126 | | | | | | WT vs. PKU | |
| | Escape latency (MWM - training day 5) | | Simple t-test | WT: 0.927<br>PKU: 0.897 | WT: p>0.05<br>PKU: p>0.05 | O | | | | | -5.226 | 16.519 | 0.0000750 | | | WT vs. PKU | |
| Supplementary Fig. 6k | Path length (MWM - training day 1) | | Simple t-test | WT: 0.914<br>PKU: 0.896 | WT: p>0.05<br>PKU: p>0.05 | O | | | | | -0.036 | 27 | 0.971 | | | WT vs. PKU | |
| | Path length (MWM - training day 2) | | Simple t-test | WT: 0.969<br>PKU: 0.969 | WT: p>0.05<br>PKU: p>0.05 | O | | | | | -3.405 | 27 | 0.00208 | | | WT vs. PKU | |
| | Path length (MWM - training day 3) | | Simple t-test | WT: 0.966<br>PKU: 0.962 | WT: p>0.05<br>PKU: p>0.05 | O | | | | | -4.768 | 27 | 0.0000570 | | | WT vs. PKU | |
| | Path length (MWM - training day 4) | | Mann-Whitney U test | WT: 0.879<br>PKU: 0.964 | WT: p<0.05<br>PKU: p>0.05 | X | 30 | 166 | -3.245 | 0.000707 | | | | | | WT vs. PKU | |
| | Path length (MWM - training day 5) | | Simple t-test | WT: 0.924<br>PKU: 0.984 | WT: p>0.05<br>PKU: p>0.05 | O | | | | | -4.005 | 27 | 0.00105 | | | WT vs. PKU | |
| Supplementary Fig. 6l | Swimming speed (MWM - probe test) | | Simple t-test | WT: 0.976<br>PKU: 0.947 | WT: p>0.05<br>PKU: p>0.05 | O | | | | | 2.968 | 27 | 0.00621 | | | WT vs. PKU | |
| Supplementary Fig. 6m | Left-quadrant occupancy (MWM - probe test) | | Simple t-test | WT: 0.922<br>PKU: 0.887 | WT: p>0.05<br>PKU: p>0.05 | O | | | | | 0.747 | 27 | 0.462 | | | WT vs. PKU | |
| | Target-quadrant occupancy (MWM - probe test) | | Simple t-test | WT: 0.918<br>PKU: 0.926 | WT: p>0.05<br>PKU: p>0.05 | O | | | | | 3.802 | 27 | 0.000746 | | | WT vs. PKU | |
| | Right-quadrant occupancy (MWM - probe test) | | Simple t-test | WT: 0.970<br>PKU: 0.942 | WT: p>0.05<br>PKU: p>0.05 | O | | | | | -0.773 | 27 | 0.446 | | | WT vs. PKU | |
| | Opposite-quadrant occupancy (MWM - probe test) | | Mann-Whitney U test | WT: 0.867<br>PKU: 0.908 | WT: p<0.05<br>PKU: p>0.05 | X | 35 | 171 | -3.026 | 0.00179 | | | | | | WT vs. PKU | |
| Supplementary Fig. 7b | OFT (total distance moved) | Ro(0 mg/kg): 7 mice<br>Ro(3 mg/kg): 7 mice<br>Ro(5 mg/kg): 6 mice | ANOVA | Ro(0 mg/kg): 0.955<br>Ro(3 mg/kg): 0.840<br>Ro(5 mg/kg): 0.890 | Ro(0 mg/kg): p>0.05<br>Ro(3 mg/kg): p>0.05<br>Ro(5 mg/kg): p>0.05 | O | | | | | | | | F(2,17)=0.053<br>p=0.949 | | | |
| Supplementary Fig. 7c | OFT (thigmotaxis) | | | Ro(0 mg/kg): 0.963<br>Ro(3 mg/kg): 0.860<br>Ro(5 mg/kg): 0.931 | Ro(0 mg/kg): p>0.05<br>Ro(3 mg/kg): p>0.05<br>Ro(5 mg/kg): p>0.05 | O | | | | | | | | F(2,17)=0.151<br>p=0.861 | | | |
| Supplementary Fig. 7e | OFT (total distance moved) | Veh: 7 mice<br>L-phe: 7 mice | Simple t-test | Veh: 0.928<br>L-phe: 0.939 | Veh: p>0.05<br>L-phe: p>0.05 | O | | | | | -0.241 | 12 | 0.814 | | | Veh vs. L-phe | |
| Supplementary Fig. 7f | OFT (thigmotaxis) | | | Veh: 0.978<br>L-phe: 0.898 | Veh: p>0.05<br>L-phe: p>0.05 | O | | | | | 0.244 | 12 | 0.812 | | | Veh vs. L-phe | |
| Supplementary Fig. 8c | Number of transients | 5 mice | Paired t-test | diff: 0.916 | diff: p>0.05 | O | | | | | -3.984 | 4 | 0.0163 | | | baseline vs. Ro | |
| Supplementary Fig. 9b | OFT (total distance moved) | WT-Veh: 10 mice<br>WT-Ifen: 10 mice<br>PKU-Veh: 9 mice<br>PKU-Ifen: 9 mice | ANOVA | WT-Veh: 0.959<br>WT-Ifen: 0.858<br>PKU-Veh: 0.947<br>PKU-Ifen: 0.982 | WT-Veh: p>0.05<br>WT-Ifen: p>0.05<br>PKU-Veh: p>0.05<br>PKU-Ifen: p>0.05 | O | | | | | | | | F(3,34)=1.412<br>p=0.256 | | | |
| Supplementary Fig. 9c | OFT (time in center) | | | WT-Veh: 0.914<br>WT-Ifen: 0.786<br>PKU-Veh: 0.929<br>PKU-Ifen: 0.965 | WT-Veh: p>0.05<br>WT-Ifen: p>0.05<br>PKU-Veh: p>0.05<br>PKU-Ifen: p>0.05 | O | | | | | | | | F(3,34)=0.337<br>p=0.799 | | | |
| Supplementary Fig. 11c | Normalized expression level | NT: 6 mice<br>KD: 6 mice | Simple t-test | NT(GluN2b): 0.881<br>KD(GluN2b): 0.943 | NT(GluN2b): p>0.05<br>KD(GluN2b): p>0.05 | O | | | | | 4.565 | 10 | 0.00103 | | | Scr vs. KD | |
| | | | | NT(GluN2a): 0.888<br>KD(GluN2a): 0.858 | NT(GluN2a): p>0.05<br>KD(GluN2a): p>0.05 | O | | | | | -0.511 | 10 | 0.621000 | | | Scr vs. KD | |
| | | | | NT(GluN1): 0.939<br>KD(GluN1): 0.878 | NT(GluN1): p>0.05<br>KD(GluN1): p>0.05 | O | | | | | -0.68 | 10 | 0.512000 | | | Scr vs. KD | |
